# Stereoselective methyl-swapping demonstrates target specificity of cognitive enhancer

**DOI:** 10.64898/2026.02.04.700910

**Authors:** Morgane Boone, Udit Dalwadi, Aniliese Deal, Ping Jun Zhu, Tristan I. Croll, Maya Yamazaki, Kimberly Prescott, Renato Minopoli, Isabella Biscocho, Junhua Wang, D. John Lee, Christopher P. Arthur, Thomas G. Laughlin, Hongyi Zhou, Matthew T. Klope, Pascal F. Egea, Michael Schoof, Rosalie Lawrence, Adam R. Renslo, Mauro Costa-Mattioli, Adam Frost, Peter Walter

## Abstract

The Integrated Stress Response (ISR) couples cellular stress sensing to translational control, playing a critical role in the homeostatic regulation of cell health. However, prolonged and unmitigated ISR activation becomes maladaptive and drives the progression of a wide range of pathologies, including cognitive decline. Pharmacological inhibition of the ISR with the small, drug-like molecule ISRIB has proven remarkably effective in reversing cognitive deficits and pathology in animal models, highlighting its potential for therapeutic intervention in humans. We engineered an allele-specific ISRIB analog (mISRIB) that selectively targets a mutant form of eIF2B, the molecular target of ISRIB, without affecting wild-type eIF2B. Notably, mISRIB treatment in mice homozygous for the eIF2B mutant allele enhances synaptic plasticity and long-term memory, confirming the on-target mechanism underlying ISRIB’s cognitive benefits. Our results provide a framework for dissecting the ISR’s contributions within complex cellular networks, such as those governing brain function, with precise temporal and spatial resolution.

## Introduction

To maintain homeostasis in the face of multiple environmental and physiological stressors, eukaryotic cells evolved various adaptive mechanisms for stress detection, mitigation, and resolution. One of these, the Integrated Stress Response (ISR), is an evolutionarily ancient signaling cascade that tunes protein synthesis through modulation of mRNA translation initiation^1,2^. By temporarily repressing general mRNA translation and simultaneously boosting the translation of a subset of uORF-containing mRNAs, the ISR diverts the cell’s energy resources towards resolution, or poises them towards apoptosis when the stress is sustained. Moreover, because some uORF-regulated mRNAs encode transcription factors, ISR activation qualitatively remodels the transcriptome. Beyond responding to stress, the ISR plays crucial roles in normal physiology, from differentiation and development^3,4^, synaptic plasticity and memory^5^, to immune system modulation^6,7^ and metabolic regulation^8–10^.

Maladaptive ISR activation, however, contributes to the pathogenesis of various – often age-related – diseases^11–16^, as well as to the cognitive decline observed with aging and brain injury^17–20^. In the past few years, pharmacological inhibition of this pathway using the small molecule ISRIB or its functional analogs demonstrated effective alleviation of neurodegenerative symptoms in an etiologically diverse array of diseases and conditions in mice and rats^21–28^. Further underscoring its potential to reverse disease in humans, chemically diverse ISRIB analogs are currently in clinical trials for Amyotrophic Lateral Sclerosis (ALS), Vanishing White Matter Disease (VWMD), and Major Depressive Disorder (MDD)^29–33^.

At the molecular level, the ISR impinges on the process of mRNA translation initiation by reducing the availability of eIF2-GTP-Met-tRNA^Meti^ ternary complex (TC), which is necessary for start codon recognition (reviewed in Hinnebusch et al, 2016)^34^. In mammals, diverse stress signals can activate one or several of the four canonical ISR stress-sensing kinases (PERK, HRI, PKR, and GCN2), all converging to the phosphorylation of the alpha subunit of trimeric eIF2^35–40^. Phosphorylation of Ser52 (historically numbered as Ser51, which omitted the cleaved-off initiating methionine) of eIF2alpha results in the conformational conversion of the eIF2 substrate into potent inhibitor (eIF2-P). eIF2 and eIF2-P exert their opposing effects via alternative binding to eIF2B, the guanine nucleotide exchange factor (GEF) that catalyzes GDP/GTP exchange on eIF2^41–45^. Hence, the phosphorylation of eIF2 by one of the ISR kinases results in the accumulation of GDP-bound eIF2 and a depletion of the functional GTP-bound eIF2 necessary for TC assembly and translation initiation.

The heterodecameric eIF2B complex (eIF2B(αβγδε)_2_) is an unusually large GEF, whose complex layers of regulation have been the subject of many recent studies and have been scrutinized up to atomic resolution. Metazoan eIF2B is allosterically regulated and oscillates between two major conformational states. The active state of the enzyme (A-State), promoted by ISRIB binding and by its substrate-bound form, adopts a ‘closed’ conformation. By contrast, binding of its inhibitor, eIF2-P, at an allosterically connected interface ‘opens’ the enzyme, reducing both substrate affinity and catalytic activity (I-State)^46,47^. We recently showed that mutations affecting the switch helix and the latch helix stabilize the enzyme at critical positions along an allosteric chain, and can thus induce or repress ISR activation in the cell in a manner that is independent of eIF2-P levels^48–50^. Similar effects can be achieved with drug-like small molecules, such as the ISR-inhibitor ISRIB. This conformational switch has been subverted by several viruses as a means of ISR evasion^50–52^, and is potentially commandeered by metabolites or other cellular factors to modulate the response.

ISRIB has been shown to be a cognitive enhancer that allosterically antagonizes eIF2-P by stabilizing eIF2B in the A-State. It binds in a distinct pocket formed at the β-δ’ and β’-δ interfaces^46,47,53,54^. In the absence of eIF2Bα, ISRIB can also substitute for the α subunit’s stapling function, bridging eIF2B(βγδε) tetramers to form a stable, catalytically active eIF2B(βγδε)_2_ octamer^53,46^. While ISRIB’s dual mechanism of action is well-understood at the atomic and molecular level, how ISRIB promotes brain health remains an open question. Motivated by the need for tools allowing precise cell-type specific, temporal control of ISR inhibition, we thought to explore the ‘bump-and-hole’ approach –akin to the orthogonal kinase-ATP analog pairs pioneered by the Shokat lab^55^ – to engineer a chemical-genetic probe based on the eIF2B::ISRIB interaction. The bumped ligand, mISRIB, is an allele-sensitive ISRIB analog with exquisite specificity for its cognate eIF2Bδ target *in vitro*, *in cellula*, and *in vivo*; while the mutated protein target retains functionality. Like ISRIB, mISRIB promotes cognitive function in mice with the cognate eIF2Bδ allele, reinforcing the on-target nature of ISRIB’s effects, and paving the way for studies dissecting interactions within complex networks, such as those governing brain function.

## Results

### A methylated ISRIB selectively promotes eIF2B-δL179V function *in vitro*

We previously noted that symmetric methylation of the methylene carbon of the ISRIB glycolamide linker severely diminishes ISRIB’s ability to inhibit the ISR in mammalian cells^53,56^. With the spatial configuration of the human eIF2B-ISRIB binding pocket in mind^53,54^, we predicted that a localized expansion of the pocket might enable to accommodate such methylated ISRIB analogs, henceforth referred to as ‘mISRIBs’. Addition of the methyl group to the methylene linker in ISRIB creates a chiral carbon that can exist as two stereoisomers, S and R. Since ISRIB is a symmetric molecule containing two glycolamide linkers, we synthesized two methylated stereoisomers mISRIB(R,R) and mISRIB(S,S) (Fig. 1A).

**Figure 1.**
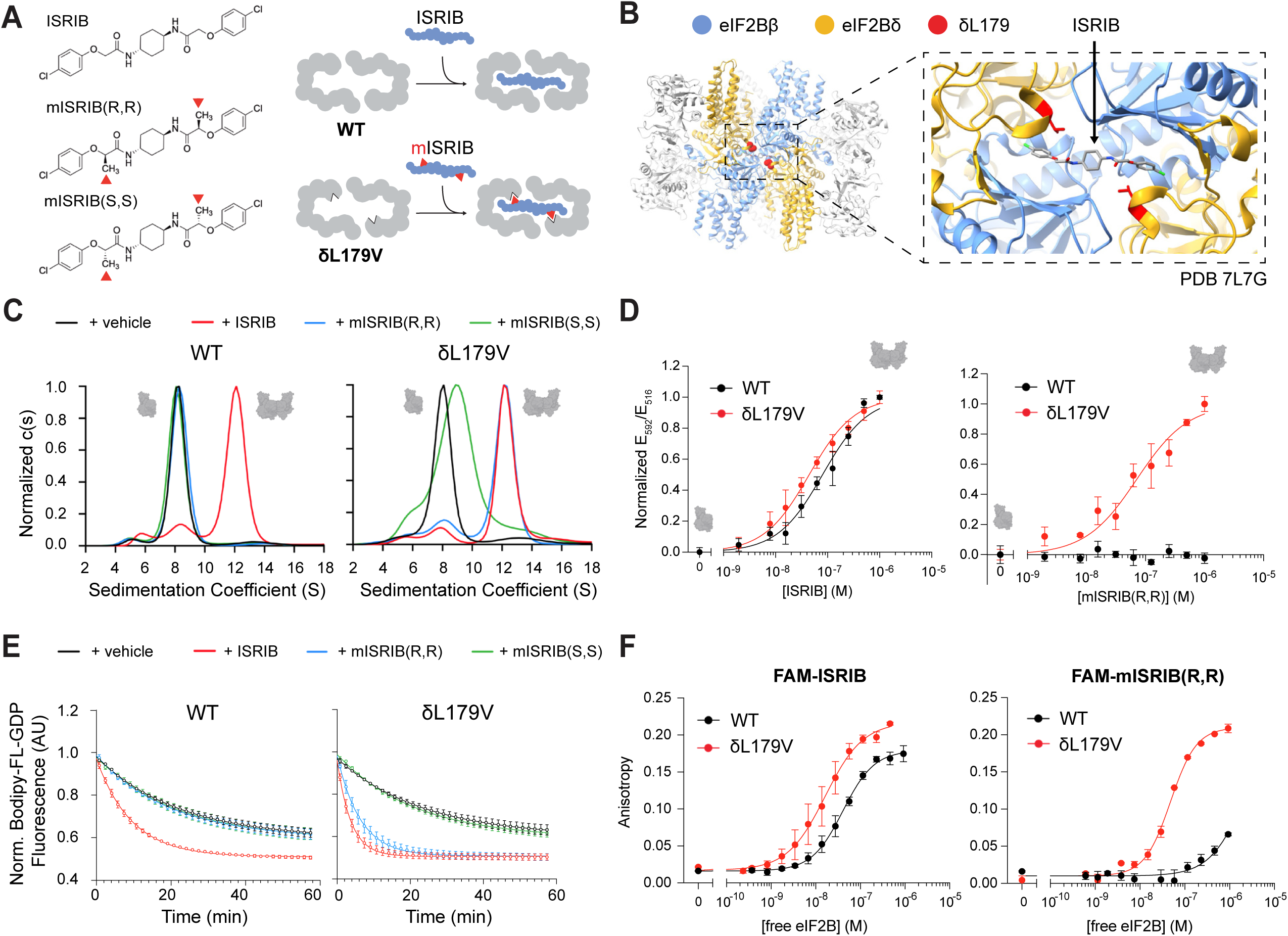
A new ligand, mISRIB(R,R), selectively binds and engages eIF2B-δL179V. (**A**) Left: methylated ISRIBs used in this study. Right: a ‘bump-and-hole’ strategy for new protein-ligand pair design. The additional methyl groups (‘bumps’, red triangles) prohibit binding of the compounds (blue) to the wild-type (WT) eIF2B ISRIB-binding pocket (grey). Targeted mutation of this binding pocket (here δL179V) can allow for the accommodation of these mISRIBs. **(B)** Position of the δL179 residue (red) in the WT eIF2B::ISRIB structure^46^ (PDB: 7L7G). **(C)** WT and δL179V eIF2B tetramer (βγδε, 1 μM) dimerization by ISRIB and mISRIB compounds (1 μM), as interrogated using analytical ultracentrifugation (sedimentation velocity), shows a selective stapling of δL179V tetramers with mISRIB(R,R). Tetramers sediment with a sedimentation coefficient of ∼8 S, octamers ((βγδε)_2_) at ∼12 S. (**D**) WT and δL179V eIF2B fluorescently-tagged tetramer (β^mNeonGreen^γδ^mScarlet-i^ε, 200 nM) dimerization by ISRIB and mISRIB compounds using FRET (1h incubation). For mISRIB(R,R), no dimerization was observed in the measured concentration range for WT tetramers, and for L179V, dimerization efficiency was similar to that of ISRIB for WT. **(E)** Guanine nucleotide exchange (GEF) activity of eIF2B tetramers (100 nM) with ISRIB and mISRIB compounds (1 μM), as assessed by BODIPY-FL-GDP unloading with unlabeled GDP on eIF2. Unloading activity of WT tetramers bound to ISRIB (t_1/2_ = 6.5 ± 0.5 min) is similar to L179V tetramers bound to mISRIB(R,R) (t_1/2_ = 4.4 ± 0.7 min), as is the intrinsic, vehicle-treated tetramer activity (t_1/2_ WT = 15.3 ± 0.3 min; t_1/2_ L179V = 16.4 ± 1.0 min). The mISRIBs, however, do not enhance WT tetramer activity. **(F)** Binding of FAM-labeled ISRIB and mISRIB(R,R) to the eIF2B WT and L179V decamer (eIF2B(αβγδε)_2_) as measured using fluorescence anisotropy. Binding affinity of FAM-ISRIB for WT decamer (*K*_D_= 38.9 ± 6.5 nM) is similar to the affinity of FAM-mISRIB(R,R):: L179V decamer (*K*_D_= 40.1 ± 6.2 nM). FAM-ISRIB binding is enhanced 2-fold by the introduction of the L179V mutation (*K*_D_= 17.3 ± 8.5 nM). **(D-F)** Biological replicates: n = 3, except for FAM-mISRIB(R,R) K_D_ in (F) where n = 2. All error bars and ‘±’ designations are s.e.m.

Amino acid L179 of the δ subunit of human eIF2B (δL179) provides one of the sidechains lining the ISRIB-binding pocket, contributing to its hydrophobicity, and playing a key role in ISRIB binding^46,57^. It is also auspiciously positioned near the ISRIB glycolamide linker carbon in the ISRIB-bound eIF2B structure (PDB 7L7G), and therefore a prime candidate for targeted mutation (Fig. 1B).

Given the need to accommodate only a single methyl group, we decided to mutate δL179 to valine*. E. coli*-expressed human eIF2B(βγδ^L179V^ε) consistently purified as a tetrameric species (henceforth referred to as “δL179V tetramers”) that was indistinguishable from wild-type (WT) eIF2B(βγδε) tetramer in both stability and sedimentation velocity analytical ultracentrifugation experiments (Fig. 1C, Suppl. Fig. 1A-B). These observations contrast with a previous report that the mutation would cause constitutive dimerization of δL179V tetramers^53^. Notably, in the presence of mISRIB(R,R), δL179V tetramers sedimented predominantly as octamers (Fig. 1C). This observation is reminiscent of ISRIB’s ability to staple WT tetramers into octamers in the absence of the eIF2B’s alpha subunit^53^. By contrast, the other mISRIB stereoisomer, mISRIB(S,S), did not fully engage δL179V tetramers at this concentration, highlighting the stereoselectivity of the mutated pocket. Wild-type eIF2B tetramers remained unaffected by either mISRIB stereoisomer, and δL179V tetramers retained the ability to be dimerized by ISRIB itself. Titration of ISRIB and mISRIB(R,R) to quantitatively measure tetramer assembly to octamers using Förster resonance energy transfer (FRET) revealed very similar octamerization behaviors (WT::ISRIB: EC_50_ = 85.7 ± 24.1 nM versus δL179V::mISRIB(R,R): EC_50_ = 82.4 ± 22.6 nM) (Fig. 1D, Suppl. Fig. 1D).

In line with these observations, measuring the guanine-nucleotide exchange (GEF) activity of WT and δL179V tetramers on Bodipy-GDP-bound eIF2 substrate similarly did not reveal any differences in absence of any drug (WT: t_1/2_= 15.3 ± 0.3 min, δL179V: t_1/2_= 16.4 ± 1.0 min), but a clear and δL179V-selective 3-fold increase in activity in the presence of mISRIB(R,R) (WT: t_1/2_= 13.6 ± 1.2 min, δL179V: t_1/2_= 4.4 ± 0.7 min) (Fig. 1E, Suppl. Fig. 1C).

Having established that the δL179V mutation does not alter the ability of eIF2B tetramers to assemble into octamers or its basal GEF activity, we next ascertained that, likewise, it would not impair the eIF2B decameric holoenzyme. Indeed, *in vitro* assembly of eIF2B(αβγδε)_2_, which is the fully active form of the complex and the predominant state of the complex in cultured K562 cells^46^, was as efficient for δL179V as for WT when measured using analytical ultracentrifugation (Suppl. Fig. 2A, C) or quantified via FRET (WT: EC_50_ = 57.1 ± 10.7 nM, δL179V: EC_50_= 76.7 ± 25.4 nM) (Fig. 2A, Suppl. Fig. 2D). Similarly, using *in vitro* pulldowns of the eIF2B holoenzyme with FLAG-tagged eIF2 immobilized on anti-FLAG beads over a range of concentrations, we did not observe any measurable differences in affinity for the substrate, eIF2 (WT: *K*_D_ = 3.5 ± 1.7 nM, δL179V: *K*_D_ = 3.0 ± 0.9 nM), or for its phosphorylated counterpart, the GEF-inhibitor eIF2-P (WT*: K*_D_ = 5.9 ± 2.5 nM, δL179V: *K*_D_ = 6.3 ± 0.6 nM) (Fig. 2B, Suppl. Fig. 2E). The nucleotide exchange activity of both mutant and WT decamer was also indistinguishable in fixed concentration GEF assays, both at baseline (WT: t_1/2_ = 7.0 ± 0.6 min, δL179V: t_1/2_ = 7.2 ± 0.8 min) and in the presence of an inhibitory concentration of eIF2-P (WT: t_1/2_ = 49.2 ± 2.2 min, δL179V: t_1/2_:= 51.1 ± 14.6 min) (Fig. 2C, Suppl. Fig. 2A-B, F-H).

**Figure 2.**
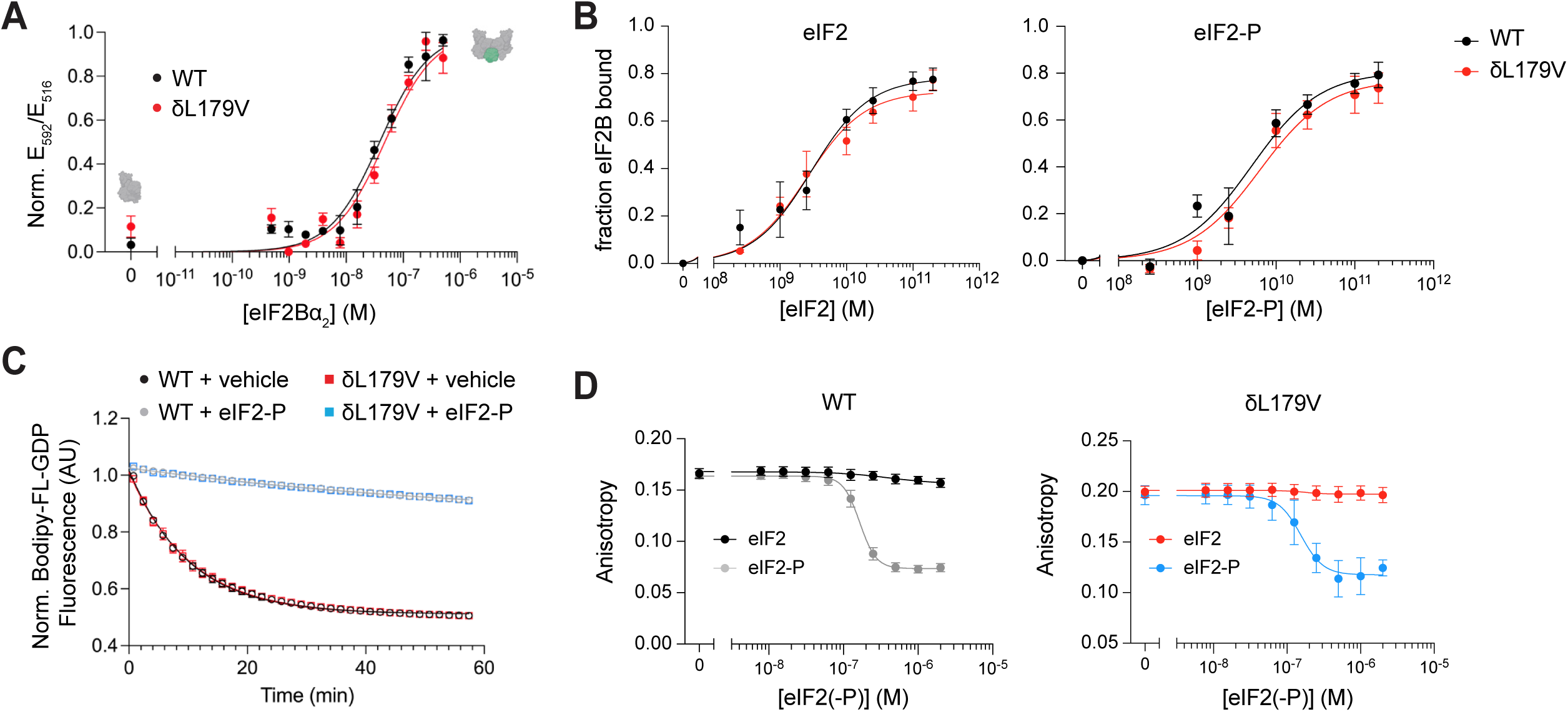
eIF2B-δL179V decamers are functionally indistinguishable from their WT counterparts, but can be allosterically modulated by mISRIB(R,R). (**A**) Holoenzyme assembly as measured by FRET after 1h of fluorescently-tagged WT or L179V eIF2B tetramer (β^mNeonGreen^γδ^mScarlet-i^ε, 50 nM, grey in inset cartoon) incubation with eIF2B(α)_2_ (green in inset cartoon)). WT EC_50_ = 57.1 ± 10.7 nM; δL179V EC_50_= 76.7 ± 25.4 nM. **(B)** Binding affinity of eIF2 or eIF2-P for WT and L179V decamers, as measured by *in vitro* pulldown with eIF2(-P) bound to magnetic beads. **(C)** Guanine nucleotide exchange (GEF) activity of WT or δL179V eIF2B decamers ((αβγδε)_2_),10 nM) with or without eIF2-P (250 nM), as assessed by BODIPY-FL-GDP unloading with unlabeled GDP on eIF2. Unloading activity of WT decamers (t_1/2_ = 7.0 ± 0.6 min) is similar to L179V decamers (t_1/2_ = 7.2 ± 0.8 min), and eIF2-P inhibits their GEF activity to the same degree (WT + eIF2-P t_1/2_ = 49.2 ± 2.2 min; L179V + eIF2-P t_1/2_ = 51.1 ± 14.6 min). **(D)** eIF2B ISRIB pocket occupancy assay as measured using fluorescence anisotropy in the presence of substrate (eIF2) or inhibitor (eIF2-P). L179V decamers are as sensitive to eIF2-P-induced FAM-mISRIB(R,R) ejection (IC_50_ = 155.4 ± 25.7 nM) as WT decamers with FAM-ISRIB (IC_50_ = 165.7 ± 12.4 nM). For (**A-D**), biological replicates: n = 3. All error bars and ‘±’ designations are s.e.m.

Fitting the initial velocities of the GEF loading reactions at various eIF2 concentrations to a Michaelis-Menten model of enzyme kinetics revealed a 25% reduction in the turnover number (WT: *k_cat_* = 20.5 ± 1.0 min^−1^, δL179V: *k_cat_* = 14.8 ± 0.8 min^−1^), but an unaffected specificity constant (WT: *k_cat_/K_M_* = 43.5 ± 3.0 min^−1^ μM^−1^, δL179V: *k_cat_/K_M_* = 46.8 ± 3.4 min^−1^ μM^−1^) (Suppl. Fig. 2J). These results suggest that the δL179V mutation results in a subtle defect in intrinsic enzymatic activity. This effect was only apparent at eIF2 concentrations more than 100-fold those of eIF2B, well above saturation binding.

We found that the results presented above for mISRIB-eIF2B tetramer interactions apply to the holoenzyme, as mISRIB(R,R) manifested a similar preference for δL179V over WT. Fluorescence anisotropy binding assays using FAM-labeled ISRIB or FAM-mISRIB(R,R) indicated very similar binding affinities of mISRIB(R,R) for δL179V decamer compared with ISRIB for WT decamer, with the *K_D_* of FAM-mISRIB(R,R)::δL179V (*K*_D_ = 43.7 ± 2.5 nM) approaching that of the FAM-ISRIB::WT complex (*K*_D_ = 42.0 ± 8.3 nM) (Fig. 1F, Suppl. Fig. 1E). FAM-mISRIB(R,R) however barely bound WT. Competing off the FAM-labeled compounds with unlabeled ISRIB, mISRIB(R,R) and mISRIB(S,S) recapitulated the findings that the mISRIBs selectively bound eIF2B-δL179V decamer ((R,R): *K*_D_= 6.8 ± 1.7 nM, (S,S): *K*_D_= 22.0 ± 5.4 nM), with little to no binding to WT within the solubility range of the compounds (Suppl. Fig. 1F-G). This selectivity was lost in a correspondingly methylated ISRIB-like molecule, the active version of the clinical ISRIB analog Fosigotifator^58^, which surprisingly bound to eIF2B WT and only displayed a ∼2-fold higher affinity for the δL179V mutant (WT: *K*_D_= 15.4 ± 2.2 nM, δL179V: *K*_D_= 6.6 ± 0.9 nM) (Suppl. Fig. 1H).

We also noticed that, aside from gaining the ability to bind mISRIBs, the δL179V mutation modestly enhanced binding of unmethylated ISRIB. For tetramers, ISRIB proved a ∼2 times more potent dimerizer of δL179V than mISRIB(R,R) (δL179V::ISRIB: EC_50_ = 48.4 ± 16.2 nM, δL179V::mISRIB(R,R): EC_50_ = 82.4 ± 22.6 nM) (Fig. 1D, Suppl. Fig. 1D). Similarly, FAM-ISRIB bound the δL179V holoenzyme with a ∼2.5-fold higher affinity than FAM-mISRIB(R,R) (FAM-ISRIB:: δL179V: *K*_D_ = 16.1 ± 7.4 nM, FAM-mISRIB(R,R):: δL179V: *K*_D_ = 43.7 ± 2.5 nM) (Fig. 1F, Suppl. Fig. 1E). All together, these observations underline that selective binding to the δL179V pocket is highly sensitive to ligand architecture.

We next verified the ability of mISRIB(R,R) to allosterically activate the eIF2B-δL179V holoenzyme. mISRIB(R,R) counteracted the inhibitory effect of eIF2-P on eIF2B δL179V’s GEF activity but was unable to do so for the wild-type eIF2B decamer (WT: t_1/2_ = 62.0 ± 14.0 min, L179V: t_1/2_ = 22.3 ± 1.8 min) (Suppl. Fig. 2F-G). Moreover, titration of eIF2-P kicked off FAM-mISRIB(R,R) from the δL179V decamer binding pocket with similar efficiency (IC_50_ = 155.4 ± 25.7 nM) as it did for FAM-ISRIB from the WT decamer (IC_50_ = 165.7 ± 12.4 nM) (Fig. 2D, Suppl. Fig. 2H). Taken together, these observations suggest that the δL179V decamer retains the ability to be allosterically regulated by eIF2-P and mISRIB through stabilization of its inhibitory I-State or active A-State, respectively.

### Cryo-EM structures reveal the molecular determinants of mISRIB selectivity

To validate the prediction that prompted the design of the methylated ISRIB, we obtained cryo-EM structures of ISRIB-bound and mISRIB(R,R)-bound δL179V holoenzymes at 2.1 Å, and 2.7 Å resolution (PDB 10GD and 10GG resp, Suppl. Fig. 5-6, Suppl. Table 1). As expected from the biochemical data above, ISRIB– and mISRIB(R,R)-bound δL179V particles exclusively adopted an active A-State conformation, characterized by (i) the ∼33 Å distance between the substrate eIF2 binding interfaces 3 and 4 (βN132-δR250), (ii) the βH160 rotamer facing the ISRIB/mISRIB binding pocket, acting as a tetramer-tetramer interface zipper, and (iii) the switch helix rotated to its active position (δR517-αD298 salt bridge) – closely resembling previously resolved A-State structures of apo-, ISRIB-bound, eIF2-bound, NSs-bound, and AcP10-bound WT eIF2B^50,51,53,54,59^ (Suppl. Fig. 3). A fourth A-State marker, the eIF2Bβ latch helix^50^, was not fully defined in either maps, similar to our previously published ISRIB-bound WT structure^53^. This observation suggests that a folded latch helix may not be strictly necessary to form an active holoenzyme in the presence of an activating ligand. Indeed, binding of ISRIB or the viral proteins NSs and AcP10 can functionally rescue latch helix deletion^50^.

As intended, the structures explain the molecular determinants of mISRIB selectivity toward δL179V (Fig. 3). In both ISRIB and mISRIB-bound δL179V decamers, similar to the ISRIB-bound WT decamer, the density in the pocket suggests that the two halves of the symmetric ligand adopt slightly different conformations due to constraints from H-bonding, hydrophobic interactions, and internal ligand geometry. To accommodate the distal chlorobenzyl groups of ISRIB, the βH188 sidechain of WT eIF2B primarily adopts an ‘outward’ rotamer, facing the center of the binding pocket (Fig. 3A). The space provided by the δL179V mutation allows ISRIB to bind within the pocket regardless of the βH188 rotamer, as shown by equal density for both rotamers (Fig. 3B). By contrast, the structure of mISRIB(R,R) bound to δL179V eIF2B reveals that the added methyl group rests directly adjacent to βH188 locked into the ‘inward’ rotamer position, and the central ring and adjacent glycolamide groups have adjusted conformations to ensure a low energy fit (Fig. 3C). Consequently, the methyl and amide groups on mISRIB sterically block βH188 from adopting the alternate rotamer. Since the ISRIB/mISRIB chlorobenzyl group requires either βH188 in the outward rotamer, or a larger space from a δL179V mutation, the WT eIF2B ISRIB pocket is rendered incompatible with mISRIB binding (Fig. 3D).

**Figure 3.**
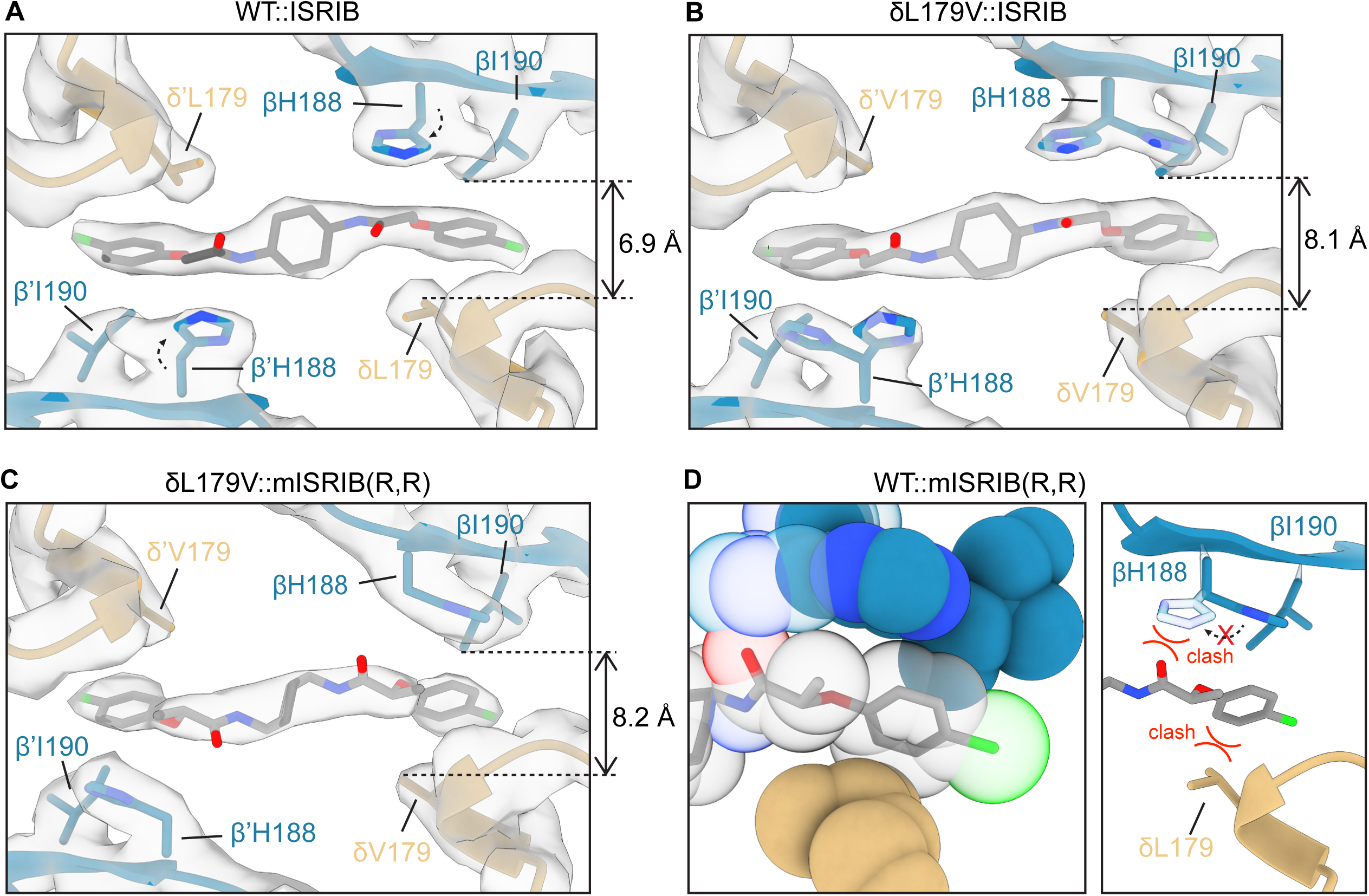
Spatial constraints in the pocket arms reveal determinants of mISRIB(R,R) binding. **(A-C)** Cryo-EM maps (grey contours) and structure models (eIF2Bβ – blue, eIF2Bδ – yellow) of indicated liganded eIF2B structures (eIF2B WT::ISRIB: PDB 7L7G), zoomed in on the pharmacophore. In WT eIF2B, ISRIB binding restricts βH188 rotamer mobility. The βH188 sidechain thus prefers the inward position (black arrow) (A). The additional space procured by mutation of δL179 to Val can counteract this, allowing βH188 to populate both orientations (B). The additional methyl group on mISRIB(R,R), however, can only be accommodated with βH188 in the inward-facing rotamer position, which necessitates more space around residue δ179 (C). **(D)** The overlay of eIF2B WT from PDB 7L7G and mISRIB(R,R) from the mISRIB-bound δL179V structure illustrates how the mISRIB oxygen would clash with the outward-facing βH188 rotamer, and how with an inward-facing βH188 and δL179, the chlorobenzene group would clash with βI190.

Following 3D classification of the apo δL179V decamer particles, we reconstructed both A– and I-State structures (PDB 10GE and 10GF resp.), although with a smaller fraction of particles adopting the A-State compared to unliganded WT eIF2B^50^ (17% δL179V vs 90% in WT) (Suppl. Fig. 4, Suppl. Table 1). While the A-State apo δL179V structure was identical to apo WT A-State, the I-State structures showed a marked loss in density in one copy of the δ subunit near the N-terminus of residue V179 (Suppl. Fig. 4D-E). The increased propensity for the I-State and the destabilization of a δ subunit may explain the slight reduction in catalytic activity in the mutant observed *in vitro* (Suppl. Fig. 2J).

In summary, mISRIB(R,R) is an allele-sensitive ISRIB analog that recapitulates ISRIB’s dual mechanism of action, but with an exclusive sensitivity for the δL179V mutant over wild-type. *In vitro*, the eIF2B decamer tolerates the δL179V mutation, as it does not measurably compromise or enhance eIF2B assembly state, stability, affinity for its substrate or its inhibitor, or strongly modulate its allosteric regulation.

### mISRIB selectively inhibits the ISR in δL179V cells

We next investigated whether this new protein-ligand pair could be used to selectively control the ISR in a cellular context. Given eIF2B-δL179V’s selectivity for mISRIB(R,R) over mISRIB(S,S), we solely focused on mISRIB(R,R) (further referred to as ‘mISRIB’) for our cellular and *in vivo* assays.

First, we CRISPR-edited two cell lines of different origin –a human cancer cell line (HEK293FlpIn-TRex) and a mouse embryonic stem cell line (AN3-12)– to introduce a homozygous L179V (L180V in mouse) mutation at the endogenous eIF2Bδ locus (*EIF2B4* in humans, *Eif2b4* in mice) (Suppl. Fig. 7). The majority of the tested clones (3/4) grew with a similar doubling time as their wild-type counterparts, suggesting that this mutation does not interfere with normal cellular growth (Fig. 4A, Suppl. Fig. 8A). The protein expression levels of various subunits of the eIF2B complex were also undisturbed in both HEKs and mouse ES cells (Fig. 4B, Suppl. Fig. 8B).

**Figure 4.**
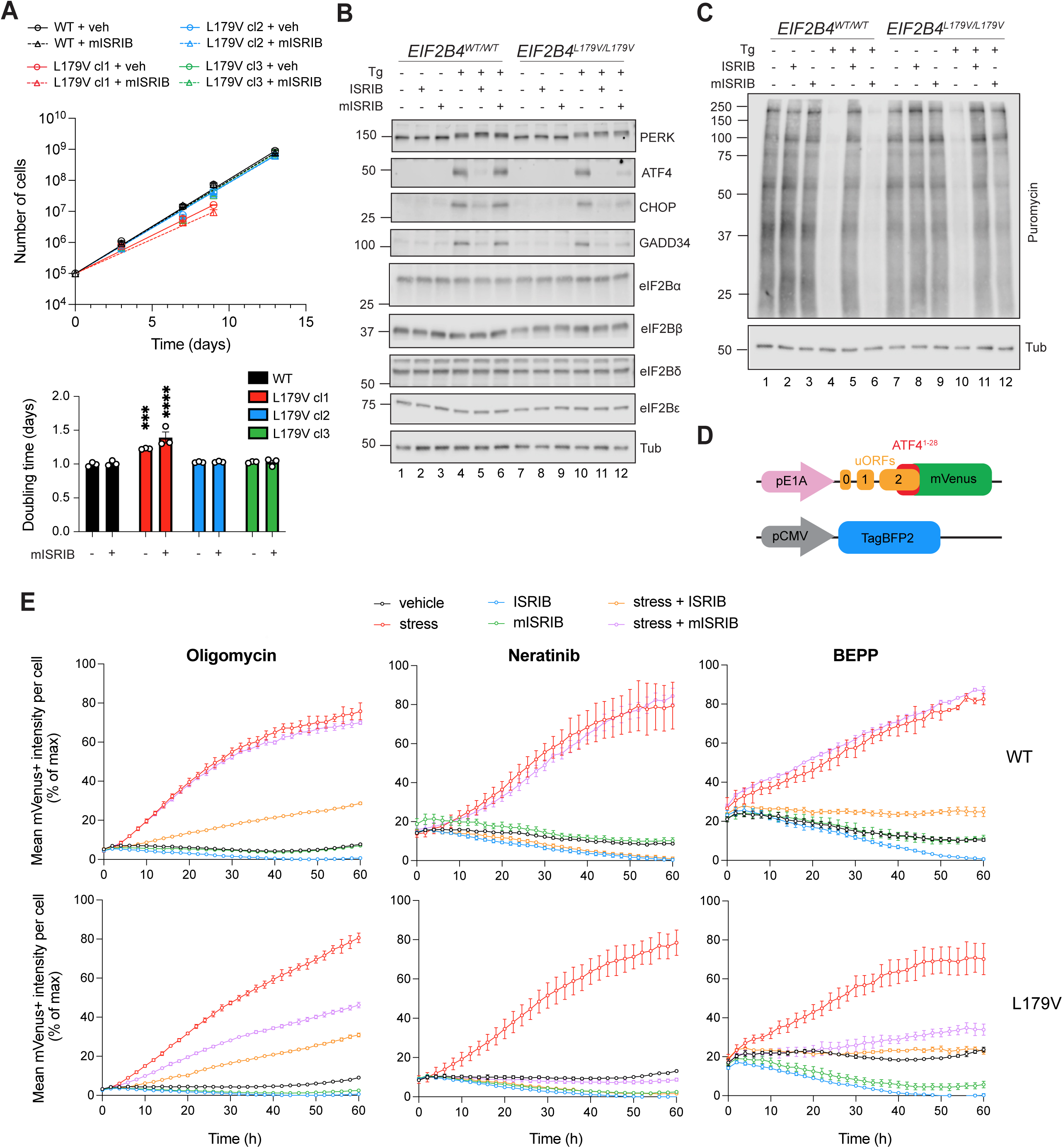
mISRIB(R,R) is a pan-ISR inhibitor in cells with eIF2B-δL179V. (**A**) Two HEK293-FlpIn T-Rex *EIF2B4^L179V/L179V^* (δL179V) clones grow at similar rates as *EIF2B4^WT/WT^*(WT) (doubling times: WT = 23.9 ± 0.7 h; δL179V cl2 = 24.6 ± 0.2 h with p=0.83, δL179V cl3 = 24.7 ± 0.3 h with p=0.83), while a third clone grows slower (δL179V cl1 = 29.4 ± 0.2 h, p=0.0004). Growth is not affected by continuous mISRIB(R,R) treatment (doubling times: WT + mISRIB = 24.2 ± 0.89 h; δL179V + mISRIB cl1= 33.4 ± 3.4 h, p<0.0001; δL179V + mISRIB cl2= 24.8 ± 0.3 h, p=0.94; δL179V + mISRIB cl3= 24.6 ± 1.3 h, p=0.98). Two-way ANOVA on doubling times, with Dunnett’s multiple comparisons vs WT. **(B)** Western blots of WT and L179V cells treated with and without stress (100 nM thapsigargin (Tg)) and/or (m)ISRIB (200 nM) for 3h. eIF2B subunit levels do not differ between cell lines. The induction of stress-induced ATF4, CHOP/DDIT3 and GADD34/PPP1R15B is repressed by mISRIB solely in a δL179V background. α-tubulin serves as a loading control. **(C)** Puromycin incorporation assay for new protein synthesis. WT and L179V cells were treated with and without stress (100 nM thapsigargin (Tg)) and/or (m)ISRIB (200 nM) for 1h, and a puromycin pulse for the last 10 mins. Stress-induced general translation initiation inhibition is repressed by mISRIB solely in the δL179V background. **(D)** ATF4 reporter cell lines were generated using a lentiviral plasmid with an ATF4 5’UTR::mVenus translation reporter and TagBFP2 selectable marker. (**E**) ATF4 translation reporter induction upon stressor treatment as measured using Incucyte live-cell imaging. mISRIB-induced ISR inhibition is genotype-specific, but independent of the nature of the stress-triggered ISR kinase. Cells were treated with 100 nM of oligomycin, 2 μM neratinib, 25 μM BEPP, and/or 200 nM ISRIB/mISRIB. For (A and E), biological replicates: n = 3. All error bars and ‘±’ designations are s.e.m. For (B-C), blots are representative of 3 biological replicates.

The edited cells furthermore retained the ability to mount an ISR response. eIF2α was phosphorylated when we acutely treated HEK293FlpIn-TRex *EIF2B4^WT/WT^*(‘WT’) and *EIF2B4^L179V/L179V^* (‘L179V’) cells with thapsigargin, a Ca^2+^-ATPase pump inhibitor that induces ER stress and subsequent activation of the PERK arm of the ISR, (Suppl. Fig. 7D, lane 4 vs 10), leading to the translational induction of proteins such as ATF4, CHOP/DDIT3, and GADD34/PPP1R15A (Fig. 4B, lane 4 vs 10) and a reduction in general protein translation, as measured by puromycin incorporation (Fig. 4C, lane 4 vs 10). As predicted from the *in vitro* work above, both WT and L179V mutant cells retain sensitivity to ISRIB (Fig. 4B, lane 5 vs 11; and Fig. 3C, lane 5 vs 11). Unlike wild-type cells, the mutation conveyed the unique ability to inhibit this response when treated with mISRIB (Fig. 4B, lane 6 vs 12; and Fig. 4C, lane 6 vs 12). These results replicated in mouse ES cells: L180V cells were indistinguishable from their wild-type counterparts in their ability to mount an ISR, but gained the specific ability to respond to mISRIB (Suppl. Fig. 8B-D). Like ISRIB, the short (3h) mISRIB treatment did not appreciably affect eIF2 phosphorylation (Suppl. Fig. 7D and 8B).

In an orthogonal assay for ISR induction, we integrated a fluorescent reporter for ATF4 mRNA translation (ATF4 5’UTR::mVenus) in the HEK293FlpIn-TRex *EIF2B4^WT/WT^* and *EIF2B4^L179V/L179V^* cells (Fig. 4D) and measured the total mVenus intensity per cell, percent cell death, and cell confluence over multiple days using live-cell imaging. Continuous mISRIB treatment suppressed ATF4 reporter induction when challenged with a broad variety of stressors (oligomycin, neratinib, and BEPP) known to target different eIF2 kinases (HRI, GCN2, and PKR, resp.)^40,60^ and counteracted the effect of these stressors on cell growth and cell death (Fig. 4D, Suppl. Fig. 9A-B). Notably, at the same 200 nM concentration, ISRIB was more potent at inhibiting ATF4 reporter accumulation in L179V cells than mISRIB, consistent with the enhanced affinity of this mutant for ISRIB *in vitro*. We obtained corresponding results when blotting for endogenous ATF4 (Suppl. Fig. 9C). Together, the experiments with different stressors underline that mISRIB activity is independent of the nature of the activated eIF2 kinase. Thus, like ISRIB, mISRIB is a true pan-ISR inhibitor. These results validate that the eIF2B-δL179V::mISRIB pair can replace eIF2B::ISRIB for physiological studies.

### mISRIB selectively enhances long-term memory *in vivo*

Genetic studies demonstrating that ISR inhibition strengthens synaptic connections and enhances long-term memory have provided a foundational understanding of the ISR’s role as a molecular rheostat for memory^5^. Building on this work, we first validated ISRIB’s ability to facilitate memory consolidation in wild-type (WT) mice using contextual fear conditioning (CFC). In this task, mice are exposed to a context (the box, i.e. conditioned stimulus) paired with an aversive foot shock (the unconditioned stimulus). Twenty-four hours later, mice are re-exposed to the same context, and freezing behavior serves as a measure of long-term memory strength. Consistent with previous results^61^, we found that daily intraperitoneal injection with ISRIB (0.25 mg/kg) for 4 days significantly enhanced freezing behavior 24h after training with a mild foot shock (1s at 0.35 mA) (Suppl. Fig. 10A), confirming that pharmacological inhibition of the ISR promotes long-term memory formation. Given that long-lasting changes in synaptic function underlie the cellular basis of long-term memory, we next measured stimulus-induced synaptic strengthening, known as long-term potentiation (LTP), in hippocampal slices from WT mice. A single high-frequency train of 100 Hz only induced a transient LTP that decayed over time. In contrast, the same protocol elicited a sustained LTP in hippocampal slices treated with ISRIB (Suppl. Fig. 10B). Thus, ISRIB treatment promotes both synaptic plasticity and long-term memory formation.

The memory and plasticity-enhancing effects of ISRIB treatment mirror those observed following genetic inhibition of the ISR^62–64^, indicating that ISRIB acts directly by ISR inhibition. However, ISRIB is a small molecule and, like all pharmacological compounds, may engage additional targets beyond eIF2B. The molecular-genetic approach developed here restricts ISRIB activity to a specific eIF2B allele and thus allowed us to address this notion with molecular precision. To this end, we generated mice homozygous for the L180V mutation in the endogenous *Eif2b4* gene (encoding eIF2Bδ) (Fig. 5A). Homozygous *Eif2b4^L180V/L180V^* mice (‘L180V mice’) were born at Mendelian ratios and were essentially indistinguishable from their age-matched wild-type counterparts: i) L180V mice exhibited comparable body weight, ii) were similarly active in an open field paradigm, iii) displayed no measurable differences in their ability to form long-term memories (Fig. 5B,D-F; Suppl. Fig. 11E). In the hippocampus and cortex, eIF2B subunit protein levels and basal ISR activity –as assessed by the abundance of eIF2-P protein and canonical ISR transcripts-was similar to WT mice (Fig. 5C, Suppl. Fig. 11A-D). Hence, the δL180V mutation does not perturb eIF2B function, ISR signaling, or normal mouse development and physiology.

**Figure 5.**
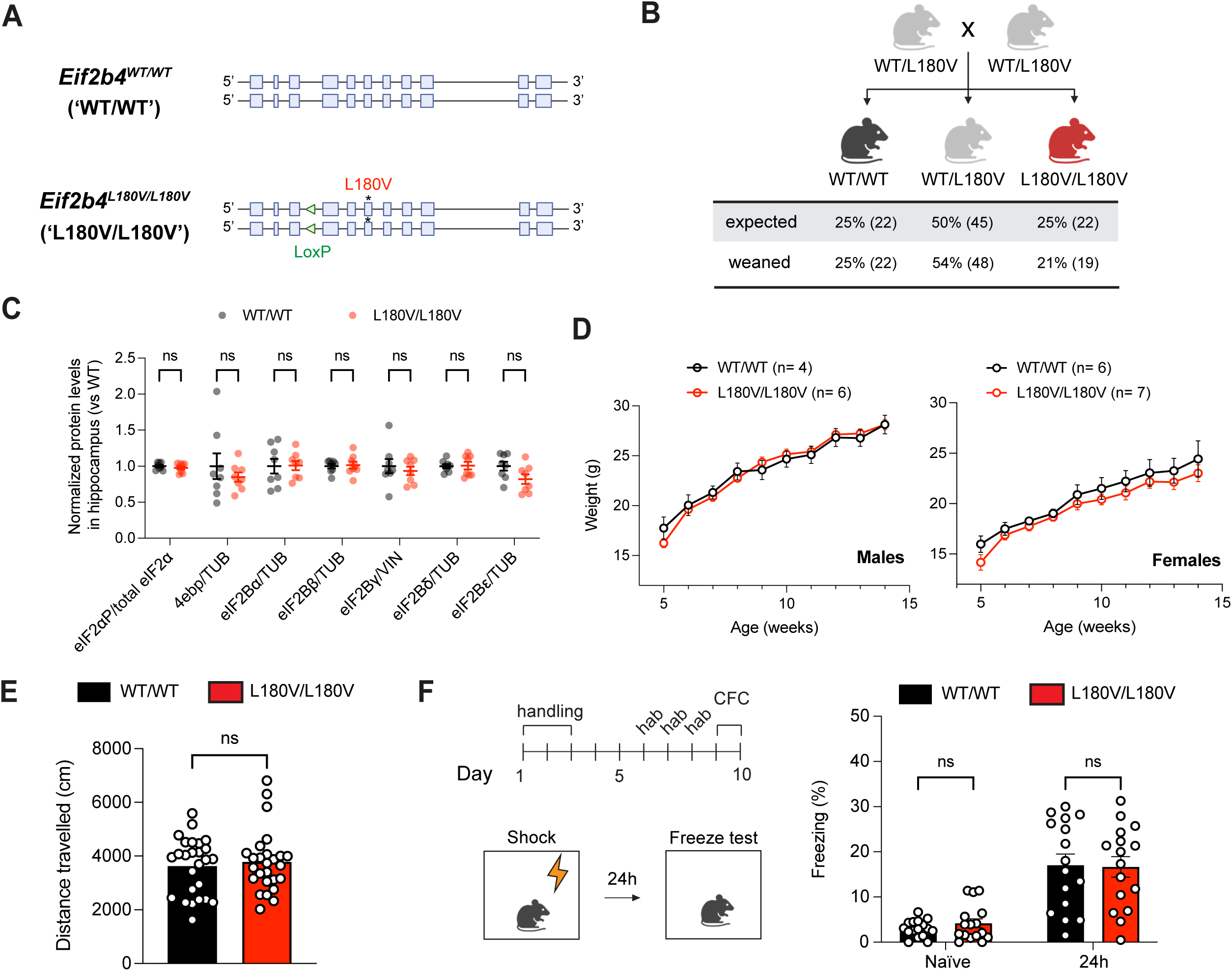
*Eif2b4^L180V/L180V^* mice are phenotypically indistinguishable from wild-type mice. (**A**) Schematic of the endogenous mouse *Eif2b4* locus in WT/WT and L180V/L180V mice. (**B**) Breeding of heterozygous *Eif2b4^WT/L180V^*(‘WT/L180V’) mice results in a Mendelian distribution of genotypes in pups born and weaned (Chi square test, Χ_2_= 0.61, p= 0.74). (**C**) Protein levels of eIF2B subunits, eIF2-P and eIF4EBP1 (‘4ebp’) in mouse hippocampus of *Eif2b4^WT/WT^* (‘WT/WT’, n= 8M) and *Eif2b4^L180V/L180V^* (‘L180V/L180V’, n= 8M) mice, as measured by western blot, and compared to wild-type as baseline. TUB= alpha-tubulin, VIN= vinculin. Two-way ANOVA with Sidak multiple comparisons test (α= 0.05). (**D**) Body weights of homozygous WT and L180V male and female mice do not differ significantly by genotype (mixed effects model, genotype *p* (males)= 0.86, genotype *p* (females)= 0.43). (**E**) Mouse locomotor activity as tracked via the average daily distance travelled in open field paradigm (WT: 12F, 14 M; L180V: 16F, 10M) (unpaired two-tailed t-test, *p*= 0.63). (**F**) In a weak training contextual fear conditioning (CFC) paradigm, L180V mice (n= 8F, 8M) do not freeze more or less than WT mice (n= 8F, 8M). Two-way repeated measures ANOVA with Sidak multiple comparisons test, naïve *p*= 0.87, 24h *p*= 0.99. For (B-F) all error bars and ‘±’ designations are s.e.m.

Like ISRIB, mISRIB crosses the blood-brain barrier following intraperitoneal injection. Using the same dosing scheme that robustly enhances long-term memory in CFC with ISRIB, we reliably detected mISRIB in plasma and brain of both WT and L180V animals 4h after the last injection at levels exceeding the *K_D_* for eIF2B-δL180V by at least an order of magnitude (Suppl. Fig. 12A). Interestingly, ISRIB accumulated at approx. 3-fold higher levels in the brains of L180V mice, perhaps due to the increased affinity of ISRIB for eIF2B-δL180V compared to WT (see Fig. 1F). mISRIB levels were substantially lower (>100-fold) in both plasma or brain 24h post-injection in either WT or L180V animals, indicating that the addition of the methyl groups on ISRIB reduces its half-life. Indeed, when incubated with mouse liver microsomes, mISRIB levels decreased by 35% within an hour (intrinsic clearance= 16.0 ± 1.0 μl/min/mg protein) whereas ISRIB remained stable (intrinsic clearance= <11.6 μl/min/mg protein) (Suppl. Fig. 12B), indicating that mISRIB metabolism by liver enzymes contributes to its reduced half-life. Based on these results, we ascertained that behavioral experiments were performed in a regime ensuring that mISRIB levels always remained well above the *K_D_* during the critical window for protein synthesis-dependent long-term memory formation (1h after training)^63–66,61^.

Based on these validations showing that the mice are normal and mISRIB enters the brain in sufficient amounts, we treated WT and L180V mice with mISRIB and assessed their long-term memory using the contextual fear conditioning paradigm described above. As expected, baseline freezing prior to training did not differ across genotypes (Fig. 6A). Rewardingly, mISRIB treatment significantly enhanced long-term memory in L180V mice, whereas WT mice remained insensitive to the compound (Fig. 6A). Consistent with these behavioral results, a single high-frequency stimulus induced a sustained LTP in mISRIB-treated hippocampal slices of L180V mice (Fig. 6B). Importantly, equally treated slices from WT mice did not produce the effect. Hence, we conclude that mISRIB recapitulates ISRIB’s memory-enhancing effects in an allelic-specific manner, directly linking the observed cognitive enhancement to eIF2B function, ruling out major contributions of off-target interactions.

**Figure 6.**
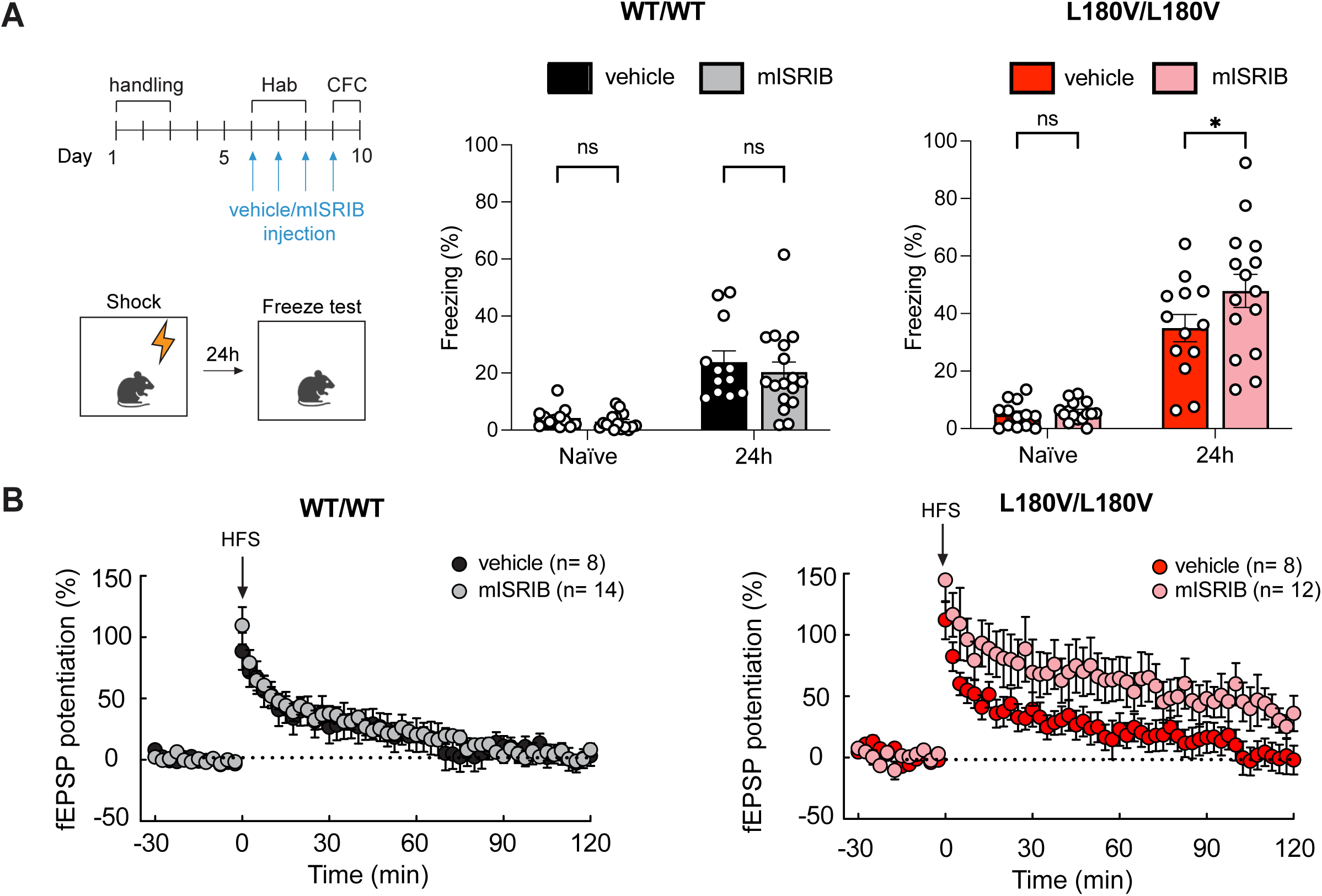
mISRIB(R,R) enhances long-term memory in constitutive *Eif2b4^L180V/L180V^* mice. (**A**) In a weak training contextual fear conditioning paradigm, daily mISRIB injection did not significantly alter freezing behavior in WT/WT mice (+vehicle n= 12, + mISRIB n= 17, male mice. Two-way repeated measures ANOVA with Sidak multiple comparisons test, naïve p= 0.93, 24h p= 0.60). In L180V/L180V mice (+ vehicle n= 13, + mISRIB n= 15), however, mISRIB injection does enhance freezing (two-way repeated measures ANOVA with Sidak multiple comparisons test, naïve p= 0.98, 24h p= 0.045). (**B**) In WT hippocampal slices, a single high-frequency pulse (HFS, 1s at 100 Hz) is insufficient to induce L-LTP, irrespective of treatment (two-tailed t-test, p= 0.33). mISRIB treatment (400 nM) only enables L-LTP induction in L180V slices (two-tailed t-test, p= 0.004). fEPSP= field excitatory post-synaptic potential. For (B-D) all error bars and ‘±’ designations are s.e.m.

## Discussion

We here describe the development and validation of mISRIB, a chemical-genetic tool for allele-specific ISR inhibition that exploits the unique pharmacophore architecture of eIF2B and its allosteric activator ISRIB. As predicted from previously obtained ISRIB-bound eIF2B structures, mISRIB’s effects are stereoisomer sensitive. Through systematic characterization spanning atomic resolution to whole-organism physiology, we demonstrated that the eIF2B-δL179V::mISRIB(R,R) mutant::drug pair enables precise, temporally-controlled ISR modulation while preserving the full mechanistic repertoire of ISRIB’s action on its cognate target.

### Definitive evidence for on-target eIF2B activation as ISRIB’s mechanism of action

Our work utilizing the eIF2B-δL179V::mISRIB(R,R) mutant::drug combination provides compelling evidence that ISRIB’s therapeutic effects are mediated exclusively through eIF2B activation and not through other potential off-target interactions. This is consistent with previous work in which mutations that interfere with ISRIB binding abolish the drug’s ability to inhibit the ISR in cells. We show that mISRIB phenocopies all measured ISRIB’s effects—from reversal of translational repression in stressed cells to enhancement of hippocampal long-term potentiation and memory consolidation—only in the presence of the mISRIB-cognate eIF2Bδ allele, thus demonstrating across resolution scales that ISRIB acts solely through eIF2B to achieve its biological effects.

### A versatile tool for dissecting ISR biology in complex cellular systems

While systemic approaches to ISR modulation have been instrumental in elucidating the pathway’s broad physiological roles, they remain fundamentally limited in their capacity to dissect the interactions occurring in complex cell assemblies, such as the brain. Tools for cell-type specific control of ISR manipulation are necessary i) to determine which cells drive specific organismal phenotypes and ii) to resolve cell-autonomous and paracrine effects of ISR modulation. Existing genetic strategies for cell-type specific ISR inhibition, including Cre-driven expression of the non-phosphorylatable eIF2α-S51A allele^65,66^ or cell-type specific promoter-driven expression of eIF2α phosphatases^67^ offer cell-type and brain region specificity, but lack temporal control. The mISRIB::eIF2B-δL179V system can address this gap, combining cell-type selectivity (achievable through conditional expression of the sensitized allele) with the temporal precision inherent to small molecule administration – akin to PKR-based chemical-genetic tools for ISR activation^68,69^.

The pharmacokinetic profile of mISRIB further enhances its utility for probing temporal dynamics of ISR function. Unlike ISRIB, which exhibits extended brain retention, mISRIB is metabolized more rapidly—levels fall close to or below the detection threshold within 24 hours of i.p. administration. This accelerated clearance, likely attributable to hepatic metabolism as suggested by our microsomal stability experiment, positions mISRIB as an ideal probe for experiments requiring discrete temporal windows of ISR inhibition. Such temporal resolution will be particularly valuable for dissecting the contribution of ISR activity during specific phases of biological processes, from the consolidation window in memory formation to developmental transitions where ISR tone must be dynamically regulated.

The observation that ISR pathway components exhibit striking cell-type-specific variation in expression levels—despite functioning as stoichiometric members of defined complexes—hints at a rich, largely unexplored layer of cell-type-specific ISR implementation^2^. The ability to selectively inhibit the ISR in defined cellular populations within intact tissue will be essential for understanding how this variation translates into functional differences and for identifying the specific cell types that drive organismal phenotypes associated with ISR modulation.

### Minimal functional perturbation of the eIF2B-δL179V allele

A critical requirement for any chemical-genetic system is that the engineered target protein retains near-native function. Our comprehensive biochemical characterization of eIF2B-δL179V reveals that the mutation preserves virtually all aspects of eIF2B function: decamer assembly, substrate and inhibitor binding affinities, and allosteric regulation by eIF2-P remain indistinguishable from wild-type. However, we did detect a subtle (∼25%) reduction in catalytic turnover (*k_cat_*), evident only under conditions where eIF2 concentrations vastly exceeded known physiological ratios to eIF2B. Importantly, the specificity constant (*k_cat_/K_M_*), which describes enzymatic efficiency under physiologically relevant substrate concentrations, was unaffected. Our cryo-EM data indicate that the apo L179V decamer retains conformational dynamics between A-State and I-State, albeit with a shift toward the I-State population – consistent with the small reduction in activity. This contrasts with other eIF2B mutations such as eIF2B-βH160D^48^, eIF2B-δL516A^49^, and eIF2B-ΔLatch^50^ where the I-State stabilization results in much larger defects in enzymatic activity. Regardless, given that cellular eIF2:eIF2B ratios are estimated at ≤10:1—far below the >100:1 ratios required to expose this kinetic defect—we anticipate negligible functional consequences under cellular conditions.

This prediction is borne out by our cell-based and organismal experiments: eIF2B-δL179V human and eIF2B-δL180V mouse cells exhibit normal growth kinetics, unperturbed ISR signaling at baseline and upon stress, and wild-type responses to ISRIB. Constitutive *Eif2b4^L180V/L180V^*mice are viable, fertile, and phenotypically normal, displaying intact hippocampal memory formation and synaptic plasticity. Nevertheless, we acknowledge that more subtle phenotypes may emerge under conditions we have not yet tested. The fact that δL179 is highly conserved across species indicates that in evolution leucine has been much preferred in this position. Even mild reductions in eIF2B GEF activity can lead to Vanishing White Matter Disease (VWMD) with advanced age^70^, indicating that eIF2B-δL180V could still function as a hypomorphic allele under metabolic or stress conditions, including aging, not recapitulated in young animals. Age-associated phenotypes or vulnerabilities to specific stressors thus remain to be systematically evaluated. Despite these caveats, the absence of overt deficits across multiple assays and life stages supports the utility of this allele for many experimental applications.

### Concluding remarks

The development of mISRIB as an allele-selective ISR inhibitor expands the experimental toolkit for dissecting this ancient and pleiotropic stress response pathway. As ISRIB analogs advance toward clinical application for neurodegenerative diseases, depression, and cognitive dysfunction, tools that enable precise interrogation of when, where, and in which cells ISR inhibition confers benefit will be essential for understanding mechanism and optimizing therapeutic strategies. The demonstration that ISRIB’s remarkable cognitive effects can be recapitulated through selective eIF2B engagement reinforces the validity of this target and provides a roadmap for future therapeutic development in the ISR space.

## Figures

**Supplementary Figure S1.** Purified eIF2B-δL179V tetramers are indistinguishable from WT, but are sensitive to mISRIB(R,R). (**A**) Coomassie-stained gel of purified tetrameric eIF2B proteins used in this study. WT and δL179V tetramers were purified from *E. coli*. (**B**) Melting temperature of WT (55.8 ± 0.2°C) and δL179V (56.2 ± 0.2°C) tetramers as measured with nano differential scanning fluorometry (nanoDSF). Unpaired two-sided t-test, *p*=0.29, n=3 biological replicates. (**C**) Half-life of Bodipy-GDP unloading of eIF2 by eIF2B tetramers and indicated compounds, n= 3 biological replicates. Two-way ANOVA with Sidak’s multiple comparison tests. WT vs L179V comparison p-values: vehicle p=0.77, + ISRIB *p*=0.01, + mISRIB(R,R) *p*<0.0001, + mISRIB(S,S) *p*=0.99. (**D**) Dimerization EC_50_s of fluorescently-tagged WT and δL179V tetramers by ISRIB (n=3 biological replicates each) and mISRIB(R,R) (n= 4 biological replicates each) using FRET. (**E**) Bar graph of *in vitro* binding *K_D_* values for FAM-(m)ISRIB compounds and WT vs δL179V eIF2B decamers, as measured using fluorescence polarization. NA: no measurable binding. (**F**) Competition of unlabeled compounds for FAM-ISRIB or FAM-mISRIB(R,R)-bound eIF2B (WT or δL179V decamers, resp.) using fluorescence anisotropy, n= 3 biological replicates. (**G**) *K_D_* values of unlabeled compounds for WT or δL179V decamers, as calculated from the fluorescence anisotropy competition experiment in (F). NA: no measurable binding. (**H**) Competition of unlabeled compounds for FAM-ISRIB or FAM-mISRIB(R,R)-bound eIF2B (WT or δL179V decamers, resp.) using fluorescence anisotropy, n= 2 biological replicates. ‘Fos’ is the active version of the prodrug Fosigotifator (ABBV-CLC-7262), ‘mFos(R,R)’ is the methylated equivalent of mISRIB(R,R). (B-H) All error bars and ‘±’ designations are s.e.m.

**Supplementary Figure S2.** The purified eIF2B-δL179V holoenzyme is functionally equivalent to the WT holoenzyme, but sensitive to mISRIB. (**A**) Coomassie-stained gel of purified decameric eIF2B and trimeric eIF2 proteins used in this study. WT and δL179V eIF2B decamers were assembled from their respective tetramers (eIF2B(βγδε)) and α-dimer (eIF2Bα_2_), both purified from *E. coli*. eIF2 trimer was purified from Expi293-F cells and phosphorylated *in vitro* by PKR. (**B**) Coomassie-stained Phostag gel of purified eIF2 and eIF2-P used in this study. Phosphorylated and non-phosphorylated eIF2α, purified from *E. coli*, are included as controls. (**C**) Decamer assembly is not affected by the δL179V mutation, as interrogated using analytical ultracentrifugation (sedimentation velocity). WT and δL179V eIF2B tetramer (βγδε, 1 μM) sediment at ∼8 S and ∼14 S, in absence or presence of eIF2B(α)_2_ (750 nM), respectively. (**D**) Dimerization EC_50_s for eIF2Bα and fluorescently-tagged WT and L179V tetramers (50 nM) as measured by FRET. Unpaired two-sided t-test, *p*= 0.52. (**E**) Binding affinities of eIF2 and eIF2-P for WT and L179V decamers. Unpaired two-sided t-tests, eIF2 binding WT vs L179V p=0.83, eIF2-P binding WT vs L179V *p*= 0.88. (**F**) Half-lives of nucleotide exchange assay using eIF2::Bodipy-FL-GDP unloading by WT and δL179V eIF2B decamers (10 nM) incubated with 1 μM indicated compounds. Two-way ANOVA with Sidak multiple comparison tests, WT vs L179V + veh *p*= 0.07, + ISRIB *p* >0.99, + mISRIB(R,R) *p*= 0.16, + mISRIB(S,S) *p*= 0.19. (**G and H**) Nucleotide exchange assay using eIF2::Bodipy-FL-GDP unloading by WT and δL179V eIF2B decamers (10 nM) incubated with indicated compounds (1 μM), with or without eIF2-P (250 nM). Statistical comparisons of exchange half-life between conditions were done using two-way ANOVA with Sidak multiple comparisons: WT vs L179V + veh *p* >0.99, + eIF2-P *p*>0.99, +eIF2-P + ISRIB *p*= 0.52, +eIF2-P + mISRIB(R,R) *p*= 0.006. (**I**) Fluorescence anisotropy IC_50_s for FAM-(m)ISRIB ejection from the eIF2B pocket by eIF2-P. Unpaired two-tailed t-test, *p*= 0.74. (**J**) Initial velocities of Bodipy-FL-GDP loading on eIF2 using WT or L179V decamer (10 nM), fit to the Michaelis-Menten equation. Unpaired two-tailed t-tests comparing WT and L179V constants; *k_cat_ p*= 0.006, *K_M_ p*= 0.06, *k_cat_/K_M_ p*= 0.49. For (C-J) all error bars and ‘±’ designations are s.e.m. Biological replicates, n=3, except for the Michaelis-Menten kinetics experiments, where n=4.

**Supplementary Figure S3.** ISRIB and mISRIB(R,R)-bound eIF2B-δL179V structures have A-State characteristics. (**A**) Snapshots of 4 allosteric sites in overlaid structure models of mISRIB(R,R)-bound δL179V decamer, ISRIB-bound δL179V decamer, A-State apo WT decamer, and I-State apo WT decamer.

**Supplementary Figure S4.** Cryo-EM analysis of apo δL179V holoenzyme. (**A**) Representative micrograph from the dataset of apo-δL179V eIF2B. (**B**) Data processing pipeline employed to generate a consensus map of δL179V eIF2B. The following analysis was done using CryoSPARC live: patch motion correction, patch CTF estimation, template picking. All subsequent steps were done using CryoSPARC v4.4+. (**C**) 3D classification of particles into 8 classes and subsequent homogeneous refinement of two separate classes produced volumes that were clearly in the A– and I-State. Distribution of particles into each class are indicated below the class images. (**D**) The final reconstruction at 2.8 Å overall resolution of A-State δL179V eIF2B (top) and a zoomed view of the map resolution and map fit surrounding the δL179V mutation. (**E**) The final reconstruction at 2.7 Å overall resolution of I-State δL179V eIF2B (top) and a zoomed view of the map resolution and map fit surrounding the δL179V mutation showing loss of density near the mutation. (**F**) Display of cryo-EM map quality and model fit of the A-State structure validating the point mutation. (**G**) Conical FSC (cFSC) summary plot generated by cryoSPARC for the A-State δL179V eIF2B reconstruction. (**H**) Conical FSC (cFSC) summary plot generated by cryoSPARC for the I-state δL179V eIF2B reconstruction.

**Supplementary Figure S5.** Cryo-EM processing of mISRIB(R,R)-bound δL179V holoenzyme. (**A**) Representative micrograph from the dataset of mISRIB-δL179V eIF2B. (**B-C**) Data processing pipeline employed to generate a consensus map of mISRIB-bound δL179V eIF2B which had poor density corresponding to the compound. The following analysis was done using CryoSPARC live: patch motion correction, patch CTF estimation, template picking. All subsequent steps were done using CryoSPARC v4.4+. (**D**) 3D classification of particles into 6 classes and subsequent homogeneous refinement of one class with density for the compound. Distribution of particles into each class are indicated below the class images. (**E**) The final reconstruction at 2.6 Å overall resolution of mISRIB(R,R)-bound δL179V eIF2B (top) and a zoomed view of the map resolution and map fit surrounding mISRIB(R,R). (**F**) Display of cryo-EM map quality surrounding mISRIB(R,R) in the final volume. (**G**) Conical FSC (cFSC) summary plot generated by cryoSPARC for the mISRIB-δL179V eIF2B reconstruction.

**Supplementary Figure S6.** Cryo-EM processing of ISRIB-bound δL179V holoenzyme. (**A**) Representative micrograph from the dataset of ISRIB-δL179V eIF2B. (**B**) Data processing pipeline employed to generate a consensus map of ISRIB-bound δL179V eIF2B which had strong compound density. The following analysis was done using CryoSPARC live: patch motion correction, patch CTF estimation, template picking. All subsequent steps were done using CryoSPARC v4.4+. (**C**) The final reconstruction at 2.1 Å overall resolution of ISRIB-bound A-State δL179V eIF2B looking at the binding pocket (left) and an alternate view after a 90° rotation (right). (**D**) Conical FSC (cFSC) summary plot generated by cryoSPARC for the ISRIB-δL179V eIF2B reconstruction. (**E**) Display of cryo-EM map quality surrounding ISRIB in the final volume.

**Supplementary Figure S7.** CRISPR-Cas9 editing of the endogenous *EIF2B4* gene with the δL179V mutation in HEK293-FlpIn-T-Rex cells. (**A**) Editing strategy at target locus of exon 6 in *EIF2B4*. The guide RNA (sgRNA) directs Cas9 for cleavage at a site close to the codon coding for L179. The provided homology-directed repair (HDR) template introduces three basepair substitutions: one for the L179V point mutation (<u>C</u>TC > <u>G</u>TC), and one silent 2 bp mutation removing a nearby BamHI restriction enzyme site (G<u>GA</u>TCC >> G<u>CT</u>TCC) to facilitate clone screening. gDNA = genomic DNA, sgRNA = single guide RNA. (**B-C**) Allele frequencies (**B**) and sequences (**C**) at the *EIF2B4* target locus in WT and δL179V cells as determined by deep sequencing. For each cell line, sequenced reads were analyzed using the CRISPResso2 pipeline. Since >99% of reads matched the HDR template in the δL179V cell line, we can infer that this cell line was homozygously edited. Unmod. = unmodified, imp. = imperfect, ambig. = ambiguous. (**D**) Western blots of *EIF2B4^WT/WT^* (WT) and *EIF2B4^L179V/L179V^*(δL179V) cells treated with and without stress (100 nM thapsigargin (Tg)) and/or (m)ISRIB (200 nM) for 3h, and probed for phosphorylated eIF2α (S51) (middle row), total eIF2α (lower row), and eIF2α on Phos-tag phospho-retention gel (upper row).

**Supplementary Figure S8.** The L180V mutation sensitizes mouse ES cells to mISRIB-induced ISR inhibition. (**A**) The L180V mutation does not measurably affect growth rate in the mouse embryonic stem (mES) cell line AN3-12. *Eif2b4^WT/WT^*(WT) and *Eif2b4^L180V/L180V^* (δL180V) cells have similar doubling times (WT = 17.6 ± 0.3 h; δL180V = 19.6 ± 0.2 h), and are unaffected by the addition of mISRIB(R,R) in the growth medium (doubling times: WT + mISRIB(R,R) = 17.8 ± 0.4 h; δL179V + mISRIB(R,R) = 19.2 ± 0.6 h). Two-way repeated measures ANOVA, effect of cell line p= 0.18 at α= 0.05. **(B-C)** Western blots of *Eif2b4^WT/WT^* (WT) and *Eif2b4^L180V/L180V^* (δL180V) cells treated with and without stress (100 nM thapsigargin (Tg)) and/or (m)ISRIB (200 nM) for 3 h, and in (B) probed for phosphorylated eIF2α (S51) (middle row), total eIF2α (lower row), and eIF2α on Phos-tag phospho-retention gel (upper row); in (C) for eIF2B subunits and proteins from ISR-responsive uORF-containing mRNAs (ATF4, CHOP/DDIT3 and GADD34/PPP1R15B). α-tubulin serves as a loading control. **(D)** Puromycin incorporation assay for new protein synthesis. Cells were treated with and without stress (100 nM thapsigargin (Tg)) and/or (m)ISRIB (200 nM) for 1 h, and a puromycin pulse for the last 10 mins. Stress-induced general translation initiation inhibition is repressed by mISRIB solely in a δL180V background.

**Supplementary Figure S9.** mISRIB is a pan-ISR inhibitor. (**A**) Cell confluence measurements with live-cell imaging in ATF4 reporter cells treated with various stressors. mISRIB treatment (200 nM) effects recapitulate the effects of ISRIB on cell growth, but in an allele-sensitive manner: with oligomycin (100 nM) and BEPP (25 μM), mISRIB hardly counteracts the stress-related growth defects, but it does so with neratinib (2 μM) stress. **(B)** Cytotox near-infrared dye cell death measurements with live-cell imaging in ATF4 reporter cells treated with various stressors. While oligomycin (100 nM) and neratinib (2 μM) do not significantly induce cell death, BEPP (25 μM) treatment does, and this is enhanced with ISRIB in WT, and ISRIB and mISRIB in L179V cells. (**C**) Western blots of *Eif2b4^WT/WT^* (WT) and *Eif2b4^L179V/L179V^* (δL179V) cells treated with and without stressors (100 nM oligomycin (Om), 1 μM neratinib (Nt), 10 μM BEPP) and/or (m)ISRIB (500 nM) for 3h, and probed for endogenous ATF4. α-tubulin serves as a loading control. For (A-B) all error bars and ‘±’ designations are s.e.m, 3 biological replicates. For western blots, blots are representative of at least 2 independent experiments.

**Supplementary Figure S10.** ISRIB enhances behavioral and cellular readouts of memory. (**A**) Male WT mice received a daily single intraperitoneal injection with vehicle (n= 14) or low-dose ISRIB (0.25 mg/kg) (n= 15) for 4 consecutive days. On the 4^th^ day of injection, 3-4h after the last injection, freezing behavior was measured for 2 mins (‘naïve’), after which mice were subjected to a single weak foot shock (0.35 mA for 1s). After 24h, ISRIB-treated mice froze significantly more than vehicle-treated mice (two-way repeated measures ANOVA with Sidak multiple comparisons test, naïve p= 0.99, 24h p= 0.0002). (**B**) L-LTP measurements induced by a single high-frequency stimulus (HFS, 1s at 100 Hz) in WT hippocampal slices treated with vehicle (n= 10) or 200 nM ISRIB (n= 7). Potentiation is significantly higher in ISRIB-treated slices when compared to vehicle-treated slices (two-tailed t-test, p= 0.024). Inset: raw traces at baseline (a) and at the end of the measurement (b). fEPSP= field excitatory post-synaptic potential. For (A-B) all error bars and ‘±’ designations are s.e.m.

**Supplementary Figure S11.** Constitutive L180V mice are similar to WT. (**A**) Western Blots of eIF2B subunits, eIF2-P and eIF4EBP1 (‘4ebp’) in mouse hippocampus of *Eif2b4^WT/WT^* (‘WT/WT’, n= 8M) and *Eif2b4^L180V/L180V^* (‘L180V/L180V’, n= 8M) mice. Control samples are from cortices of mice with and without a genetically-encoded persistent ISR activation (Crep(RR) = wild-type aka *Ppp1r15b^WT^, Crep(CC) =* mutant aka *Ppp1r15b^R658C^*). Tub= alpha-tubulin. (**B**) As in (A), but from the cortex. (**C**) Quantification of cortical protein Western Blots from (B). Two-way ANOVA with Sidak multiple comparisons test (α= 0.05). (**D**) Compared to the cortex of age-matched mice with genetically encoded persistent ISR activation (Crep(RR) = wild-type aka *Ppp1r15b^WT^, Crep(CC) =* mutant aka *Ppp1r15b^R658C^*) (n= 3 per group), transcript levels of ISR genes in the cortex of L180V mice (n= 8 per group) are not significantly elevated compared to WT mice. Stars (*) indicate genes with a min. 25% change (log2(foldchange)>|0.32|) in transcript levels in the mutant vs WT mice, with an adjusted p-value < 0.05. (**E**) Setup for strong training Contextual Fear Conditioning (CFC) experiment with WT and L180V mice. Hab= habituation. We used a CFC paradigm with strong training to test for potential long-term memory deficiencies. Freezing behavior was measured for 2 mins (‘naïve’), after which mice were subjected to two foot shocks (0.7 mA for 2s, 30s in between). After 24h, L180V mice (n= 16F, 10M) did not freeze significantly less than WT mice (n= 12F, 14M) (two-way repeated measures ANOVA with Sidak multiple comparisons test, naïve *p* >0.99, 24h *p* >0.99). For (C, E) all error bars and ‘±’ designations are s.e.m.

**Supplementary Figure S12.** mISRIB pharmacokinetics. (**A**) Total ISRIB and mISRIB concentrations in mouse plasma and brain cortex, after a once daily dosing for 4 days and tissue isolation 4h or 24h after the last injection. (**B**) Compounds were incubated with mouse liver microsomes in triplicate, and total concentrations were measured at indicated timepoints using LC-MS. For (A-B) all error bars and ‘±’ designations are s.e.m.

Table 1.

**Table 1**
| **Antibody target** | **Dilution** | **Manufacturer** | **Cat. number** |
| --- | --- | --- | --- |
| eIF2α (total) | 1/1000 | Cell Signaling Technologies | 5324S |
| eIF2α-P | 1/1000 | Cell Signaling Technologies | 9721S |
| eIF2α-P | 1/1000 | Cell Signaling Technologies | 3398 |
| eIF2α-P | 1/1000 | Cell Signaling Technologies | 3597S |
| eIF2Bα | 1/1000 | ProteinTech | 18010-1-AP |
| eIF2Bβ | 1/1000 | ProteinTech | 11034-1-AP |
| eIF2Bγ | 1/1000 | ProteinTech | 11296-2-AP |
| eIF2Bδ | 1/1000 | ProteinTech | 11332-1-AP |
| eIF2Bε | 1/1000 | ProteinTech | 29935-1-AP |
| eIF2Bε | 1/1000 | Santa Cruz | sc-55558 |
| ATF4 | 1/1000 | Cell Signaling Technologies | 11815S |
| GADD34 | 1/1000 | ProteinTech | 10449-1-AP |
| CHOP | 1/1000 | Cell Signaling Technologies | 2895S |
| α-Tubulin | 1/1000 | Cell Signaling Technologies | 3873T |
| Tubulin-Rhodamine | 1/1000 | BioRad | 12004166 |
| Vinculin | 1/1000 | Cell Signaling Technologies | 13901 |
| eIF4EBP1 | 1/1000 | Cell Signaling Technologies | 9644S |
| eIF4EBP1-P | 1/1000 | Cell Signaling Technologies | 2855 |
| Puromycin | 1/10000 | Millipore | MABE343 |

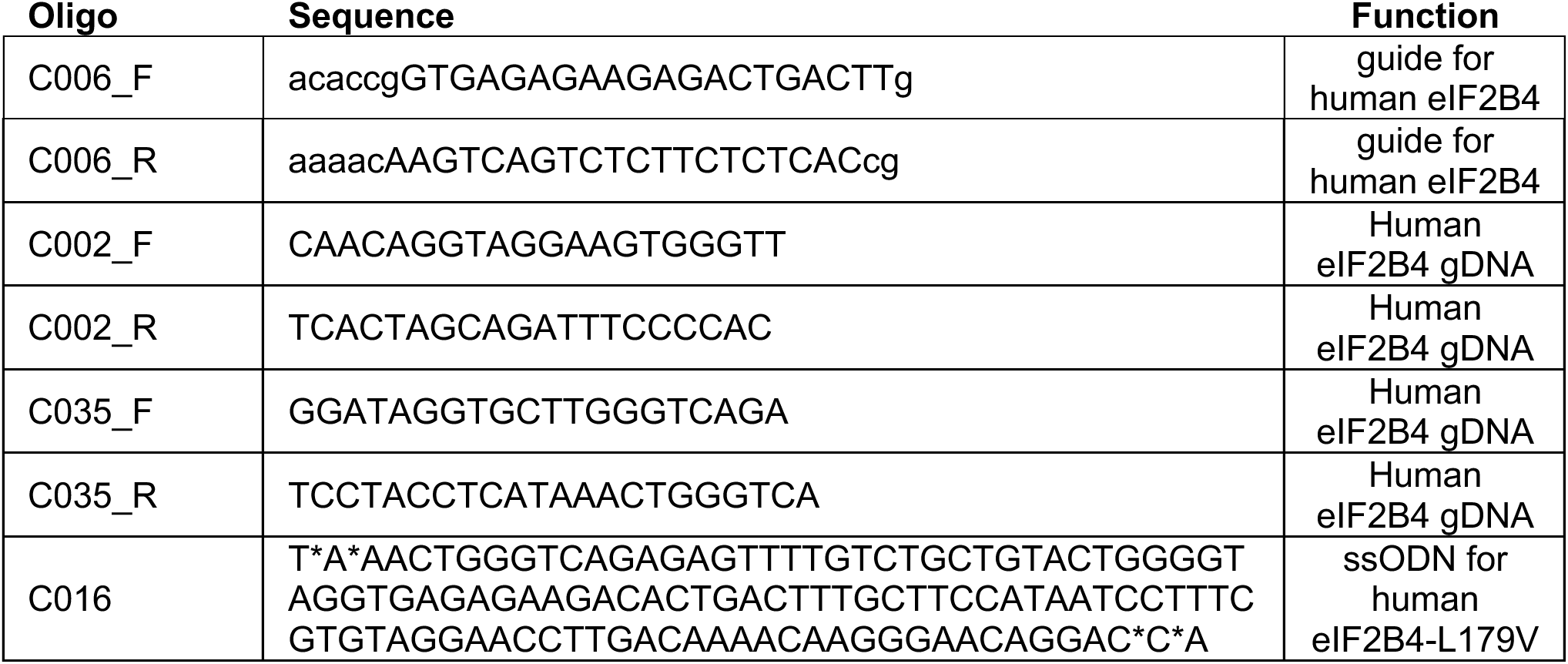
Table 2.

Table 3.

**Table 3**
| **Compound** | **Source** | **Identifier** |
| --- | --- | --- |
| ISRIB | In-house synthesis | Tsai et al, 2018 |
| ISRIB | Fisher Scientific | 5284 |
| mISRIB(R,R) | In-house synthesis | Tsai et al, 2018 |
| mISRIB(R,R) | Acme Bioscience | Custom synthesis |
| mISRIB(S,S) | In-house synthesis | Tsai et al, 2018 |
| mISRIB(S,S) | Acme Bioscience | Custom synthesis |
| FAM-ISRIB | Merken Biotech | Custom synthesis |
| FAM-mISRIB(R,R) | Merken Biotech | Custom synthesis |
| Thapsigargin | Sigma-Aldrich | SML1845-4MG |
| Oligomycin | Sigma-Aldrich | 75351-5MG |
| Neratinib | MedChemExpress | HY-32721 |
| BEPP monohydrochloride | Sigma-Aldrich | B2938-5MG |
| Puromycin dihydrochloride | Fisher Scientific | A1113803 |

## Online Methods

### Cloning

For CRISPR editing of the *EIF2B4* gene in HEK293-Flp-In-T-rex cells, guide RNAs were designed using the Benchling CRISPR gRNA Design Tool, selecting the guide with the best on-target and off-target scores, and the L179V mutation within 10 bp of the cut site. Cloning of the guide into the guide expression plasmid (MLM3636, with human U6 promoter) was done as previously described (Kwart et al. 2017). In brief, the guide RNA sequence was synthesized as single stranded DNA oligos (C006_F and C006_R) that were first annealed at 2 µM in 1x annealing buffer (40 mM Tris-HCl pH 8.0, 20 mM MgCl_2_, 50 mM NaCl, 1 mM EDTA pH 8.0), for 5 min at 95°C followed by gradual decrease of – 0.1°C/s to 25°C. The MLM3636 plasmid was digested using BsmBI (NEB) in NEB Buffer 3.1 for 2h at 55°C, and the 2.2 kb backbone was isolated from a 0.8% agarose gel with 1x SYBR Safe, and purified using the NucleoSpin Gel and PCR cleanup kit (Macherey Nagel). Backbone and annealed guide template were ligated for 1h at room temperature using T4 DNA Ligase (NEB), 100 ng backbone, 200 nM guide template, and 1x T4 DNA Ligase buffer (NEB), prior to transformation to *E. coli*. Plasmid was sequence verified by Sanger sequencing.

To generate a FRET-compatible eIF2B(βγδε) tetramers with δL179V mutation, the WT eIF2B4 sequence in pMS029^46^ was replaced with the eIF2B4-L179V sequence from pJT090^53^ using NdeI (NEB) and SbfI-HF (NEB) restriction enzyme digestion, agarose gel purification, and ligation. The resulting plasmid, pMB001, was sequence verified by Sanger sequencing.

For the expression of human trimeric eIF2, eukaryotic expression plasmids pALT002 (full-length eIF2α), pALT003 (FLAG-TEV-eIF2β), and pALT004 (full-length eIF2γ) were constructed and sequenced by Twist Bioscience.

The lentiviral ATF4-mVenus reporter plasmid (pMB033) was constructed by Vectorbuilder.

### Purification of human eIF2B tetrameric subcomplexes

WT eIF2B(βγδε) (pJT073 and pJT074 co-expression), eIF2B(βγδ^L179V^ε) (pJT090 and pJT074), WT eIF2B(β^mNeonGreen^γδ^mScarlet-i^ε) (pMS029 and pJT074 co-expression), and δL179V eIF2B(β^mNeonGreen^γδ^mScarlet-i^ε) (pMB001 and pJT074 co-expression) were expressed in *E. coli* purified over a His affinity column, anion exchange, and size exclusion as previously described^46,53^.

### Purification of human eIF2B(α)_2_ dimers

We initially used His-tagged eIF2B(α)_2_ or His-Strep-tagged eIF2B(α)_2,_ and later switched to untagged eIF2B(α)_2_ to boost the yield of the final protein. All versions gave the same results in GEF and assembly assays and are considered equivalent.

For purification of His-tagged eIF2B(α)_2_ (pJT075) dimers and His-Strep-tagged eIF2B(α)_2_ (pMB019) dimers, transformed *E. coli* BL21*-DE3 cells were grown up to OD_600_ of 0.6 in the appropriate antibiotics at 37C, and induced with IPTG for 16h at 16°C. Cells were pelleted and resuspended in ice-cold lysis buffer (20 mM HEPES-KOH pH 7.5, 250 mM KCl, 5 mM MgCl_2_, 1 mM TCEP, 20 mM imidazole, 1x Complete no EDTA protease inhibitor), and lysed by triple passage through the Emulsiflex high-pressure homogenizer (Avestin). Supernatans was clarified by ultracentrifugation for 30 mins at 4°C and 30,000x g. Subsequent purification steps were conducted on the ÄKTA Pure (GE Healthcare) system at 4°C. The lysate was loaded onto a 5-mL Histrap HP column, and protein was eluted with a linear gradient of a high-imidazole buffer (20 mM HEPES-KOH pH 7.5, 30 mM KCl, 5 mM MgCl_2_, 1 mM TCEP, 300 mM imidazole).

For pJT075 (His-eIF2B(α)_2_), the HisTrap elution was then passed through an anion exchanger MonoQ HR 10/100 GL in a low salt buffer (20 mM HEPES-KOH pH 7.5, 30 mM KCl, 5 mM MgCl2, 1 mM TCEP), and the flowthrough was concentrated with a 30 kDa mass cutoff Amicon Ultra-15 concentrator (EMD Millipore) prior to overnight dialysis in a high pH buffer (20 mM HEPES-KOH pH 8.25, 30 mM KCl, 5 mM MgCl2, 1 mM TCEP). This dialysed sample was then run over a HiTrap HP column in a high pH, low salt buffer (20 mM HEPES-KOH pH 8.25, 30 mM KCl, 5 mM MgCl2, 1 mM TCEP), and the flowthrough was once more concentrated over an Amicon column. Finally, the sample was passed over a size exclusion column (S200 Increase 10/300 GL (Cytiva)) equilibrated in S200 buffer (20 mM HEPES-KOH, pH 7.5, 200 mM KCl, 1 mM TCEP, 5 mM MgCl_2_, 5% glycerol).

For pMB019 (His-Strep-eIF2B(α)_2_), after elution from the HisTrap HP column, fractions with eIF2Bα protein were further affinity purified on a StrepTag HP column, with a step gradient using desthiobiotin (20 mM HEPES-KOH, pH 7.5, 30 mM KCl, 1 mM TCEP, 5 mM MgCl2, 2.5 mM desthiobiotin). The relevant elution fractions were then combined and concentration, and then further polished off on the S200 size exclusion column as above. For untagged eIF2B(α)_2_, dimers were purified as described previously^50^.

All proteins were quality-checked on SDS-PAGE and Western Blots for purity, stoichiometry, and identity verification, and stored aliquoted at –80°C until use.

### Purification of heterotrimeric human eIF2 and eIF2-P

Human eIF2 was purified from Expi293-F cells by Viva Biotech, as previously described^50^. Briefly, eIF2 plasmids (pALT002, pALT003, pALT004) were co-expressed with an expression plasmid for Cdc123 (pALT005) in 500 ml Expi293-F cells. Equimolar ratios (0.75 μg of each plasmid per mL of cell culture) were diluted in Opti-MEM media (Gibco) to a final concentration of 18 μg/mL, and combined with an equivalent volume of PEI at 0.12 mg/mL in Opti-MEM. The mixture was added dropwise to cells at a density of ∼3 million cells/mL to transfect the plasmids. The cells were grown for 20 hours on an orbital shaker at 37 °C, 8% CO_2_ at a speed of 125 rpm before addition of Expi Enhancer to the culture. Cell pellets were harvested by centrifugation 72 h after transfection. Pellets were resuspended with 5 mL/g lysis buffer (20 mM HEPES KOH pH 7.4, 300 mM KCl, 1 mM MgCl_2_, 1 uM Triton X-100, 1x cOmplete EDTA-free protease inhibitor cocktail [Roche]) and lysed by sonication (3 s on, 5 s off, 200 W for 100 cycles). Lysates were clarified by two consecutive centrifugations for 30 mins at 20300 x g, at 4 °C. The supernatant was incubated with anti-FLAG M2 affinity gel (Sigma) and bound protein was eluted with 100 μg/ml of FLAG peptide. GDP was added to the eluate at 1 mM and incubated for one hour. For phosphorylation, GST-PKR was added to the solution at 1:20 (PKR:: eIF2, w/w) and ATP was added to a final concentration of 1 mM. The protein, treated with/without PKR kinase, was finally run on a size exclusion column (Superdex 200 Increase 10/300 GL (Cytiva)) with S200 buffer (50 mM HEPES-KOH pH 7.5, 100 mM KCl, 2 mM MgCl_2_, 0.5 mM TCEP, 5% glycerol), concentrated by ultra-filtration (Amicon, MWCO 30 kD) and stored aliquoted at –80°C until use. Quality of the trimer was confirmed by SDS-PAGE (stoichiometry and purity check), Phostag gel (phosphorylation status), mass spectrometry (modifications and identity), and GEF assay (activity, see below).

### Analytical ultracentrifugation (AUC)

Analytical ultracentrifugation (sedimentation velocity) experiments were performed as previously described using the ProteomeLab XL-I system (Beckman Coulter) (Tsai et al. 2018). In brief, samples were loaded into cells in a buffer of 20 mM HEPES-KOH pH 7.5, 150 mM KCl, 1 mM TCEP, and 5 mM MgCl_2_. A buffer-only reference control was also loaded. Samples were then centrifuged in an AN-50 Ti rotor at 40,000 rpm at 20°C and 280 nm absorbance was monitored. Subsequent data analysis was conducted with Sedfit using a non-model-based continuous c(s) distribution.

### Nano-Differential Scanning Fluorimetry (NanoDSF)

Protein stability assays were performed by MedChemExpress (Mountain Junction, NJ), measured on a Prometheus NT.48 protein stability analysis system (Nanotemper) with standard 24-capillary chips. Each capillary was loaded with 5 µL of protein sample (final 1 µM) and 5 µL of PBS. Fluorescence measurements were performed on individual capillaries at 20 °C using the High Sensitivity mode (10 acquisitions, 5 s each). A temperature ramp from 20 °C to 95 °C was applied at a rate of 5 °C/min. Excitation power was set to 20%. The first derivative of the fluorescence ratio was used to determine the protein melting temperature.

### *In vitro* FRET assays

*In vitro* equilibrium FRET assays were performed as previously described (Schoof et al. 2021) with minor modifications. Briefly, 20 μl reactions were set up in triplicates with 50 or 200 nM WT or L179V eIF2B(β^mNeonGreen^γδ^mScarlet-i^ε) + a serial dilution titration of ISRIB, mISRIB(R,R) or eIF2B(α)_2_ in FP buffer (20 mM HEPES-KOH pH 7.5, 100 mM KCl, 5 mM MgCl2, 1 mM TCEP), in 384 square-well black-walled, clear-bottom polystyrene assay plates (Corning). Final content of NMP was 0.5% across. After 1h of incubation at 23°C, measurements were taken using the ClarioStar PLUS plate reader (BMG LabTech). mNeonGreen was excited (470 nm, 16 nm bandwidth), and mNeonGreen (516 nm, 8 nm bandwidth) and mScarlet-i (592 nm, 8 nm bandwidth) emission were monitored. FRET signal (E_592_/E_516_) was calculated as the ratio of mScarlet-i emission after mNeonGreen excitation and mNeonGreen emission after mNeonGreen excitation, and this was further normalized to the min and max signal per protein to correct for differences in brightness due to prep-to-prep variation in photobleaching. Experiments were repeated 3 to 4 independent times. Data were plotted in GraphPad Prism 10, and curves were fit to a sigmoidal curve according to the function y=1/(EC_50_ + x), assuming a Hill slope of 1.

### Guanine nucleotide exchange (GEF) assay

*In vitro* detection of GDP binding to eIF2 was performed as described previously ^48^: purified human eIF2 (100 nM) was incubated with 100 nM BODIPY-FL-GDP (Thermo Fisher Scientific) in assay buffer (20 mM HEPES-KOH, pH 7.5, 100 mM KCl, 5 mM MgCl_2_, 1 mM TCEP, and 1 mg/ml BSA) to a volume of 18 µl in 384 square-well black-walled, clear-bottom polystyrene assay plates (Corning). For the assay buffer, TCEP and BSA were always freshly added the day of the experiment. For the tetramer GEF assays, a 10X GEF mix was prepared containing 1 µM eIF2B(βγδε) tetramer (WT or δL179V), 2% N-methyl-2-pyrrolidone (NMP), and with or without 10 µM ISRIB, again in assay buffer. For the assay, 2 µl of the 10x GEF mix was spiked into the eIF2::BODIPY-FL-GDP mix, bringing the final concentrations to 100 nM tetramer, 0.2% NMP and with or without 1 µM compound. Fluorescence intensity was recorded every 10 s for 40 s prior to the 10X GEF mix spike, and after the spike for 60 min, using a Clariostar PLUS (BMG LabTech) plate reader (excitation wavelength: 477 nm, bandwidth 14 nm; emission wavelength: 525 nm, bandwidth: 30 nm). Immediately after the loading assay, in the same wells, we spiked in unlabeled GDP to 1 mM to measure unloading, again recording fluorescence intensities every 10s for 60 min as before. For clarity, datapoints were normalized to the baseline, averaged at 1 min intervals and then plotted as single datapoints. For the loading assay, data was fit to the exponential equation Y=Y_0_ + (Plateau-Y_0_)*(1-exp(-K*x)) (one-phase association) with Y_0_ set to 0. For the unloading assay, data was fit to the exponential equation Y=(Y_0_ – Plateau)*exp(-K*X) + Plateau (one-phase decay).

For assays with eIF2B(αβγδε)_2_ decamers (WT or δL179V), decamers were first assembled by combining eIF2B(βγδε) tetramer (WT or δL179V) with eIF2Bα_2_ dimer in a 2:1.5 molar ratio (a 1.5-fold excess of eIF2Bα_2_ dimer compared to the number of eIF2B(βγδε)_2_ octamers) at room temperature for 45 min. The 10X GEF mix for decamer assays contained 100 nM eIF2B(αβγδε)_2_ decamer (WT or δL179V) in assay buffer, for a final of 10 nM decamer. The ensuing steps were performed as described for the GEF assays with tetramers.

### Michaelis-Menten kinetics

Michaelis-Menten kinetic analysis of eIF2B(αβγδε)_2_ decamers (WT or δL179V) GEF activity was performed as described previously)^48^. Briefly,BODIPY-FL-GDP loading assays were performed as described above, keeping final decamer concentrations at 10 nM, but varying substrate concentration from 0 to 3 μM. BODIPY-FL-GDP concentration was kept at 2 μM final. The initial velocity (*v_0_*) was determined by a linear fit to time points acquired at 5s intervals from 50 to 200 s after addition of decamer. To convert fluorescence intensities to pmol substrate, the gain in signal after 60 min was plotted against eIF2 concentration for the 37 nM –1 μM concentrations. *Vmax* and *K_M_* were determined by fitting the initial velocities as a function of eIF2 concentration to the Michaelis–Menten equation in GraphPad Prism 10. For statistical comparisons of *Vmax, k_cat_* and *K_M_*, we used a two-sided t-test with α = 0.05, comparing each constant derived from the individual fit of each replicate experiment.

### Fluorescence anisotropy assays

For measurement of FAM-(m)ISRIB K_D_ for eIF2B(αβγδε)_2_ decamers (WT or δL179V), reactions were set up according to the protocol by Rossi et al^71^. First, decamers were assembled by combining eIF2B(βγδε) tetramer (WT or δL179V) with eIF2Bα_2_ dimer in a 2:1.5 molar ratio (a 1.5-fold excess of eIF2B(α)_2_ dimer compared to the number of eIF2B(βγδε)_2_ octamers) at room temperature for 1h, and then diluted in FP buffer (20 mM HEPES-KOH pH 7.5, 100 mM KCl, 5 mM MgCl_2_, 1 mM TCEP) in a 2-fold dilution series. FAM-ISRIB or FAM-mISRIB(R,R) were first diluted to 2.5 μM in 100% NMP prior to dilution to 50 nM in 2% NMP (always in black-walled Eppendorf tubes). Reactions were assembled in 20 ul in 384-well non-stick black plates (Corning 3820) using the ClarioStar PLUS (BMG LabTech) at room temperature, with final concentrations of 2.5 nM FAM-labeled compound, 0.5% NMP, and varying eIF2B decamer concentrations. To measure the non-specific signal, an extra row was included with an additional 1 uM of unlabeled ISRIB or mISRIB(R,R). A ‘FAM-(m)ISRIB-only’ control was included to determine gain and polarization baseline, and a ‘protein-only’ control was included to measure background polarization. The plate was covered, shaken for 1 min, spun down and allowed to incubate at 23°C for 30 mins prior to measurement of parallel and perpendicular intensities (excitation: 482 nm; emission: 530 nm). Specific anisotropy was plotted against free eIF2B concentration, fitted in GraphPad Prism 10 to a 4 parameter logistic equation Y=Bottom + (Top-Bottom)/(1+10^(LogEC^_50_^−X)*HillSlope^)), and K_D_ was calculated as the EC_50_-(FAM-(m)ISRIB/2).

For the competition assays, 20 μl reactions were set up as in Schoof et al.^46^ with 100 nM eIF2B(αβγδε)_2_ decamer (WT or δL179V), 2.5 nM FAM-(m)ISRIB in FP buffer, and a dilution series of indicated compounds in DMSO. Final DMSO concentration was 2.5%. Controls included a FAM-(m)ISRIB only reaction, a protein-only reaction, and a 300 nM decamer reaction with 2.5 nM FAM-(m)ISRIB as a max signal control. Reactions were incubated at 23°C for 30 min prior to measurement of parallel and perpendicular intensities (excitation: 482 nm; emission: 530 nm). Specific anisotropy signal and unlabeled compound K_D_’s were calculated as in Rossi et al. Data were plotted in GraphPad Prism 10, and where appropriate, curves were fit to a 4 parameter logistic equation Y= Bottom + (Top-Bottom)/(1+(IC_50_/X)^HillSlope^)).

For the pocket occupancy assays with eIF2 and eIF2-P, 20 μl reactions were set up as in Schoof et al.^46^ with 100 nM eIF2B(αβγδε)_2_ decamer (WT or δL179V), 2.5 nM FAM-(m)ISRIB in FP buffer, and a dilution series of eIF2 or eIF2-P. Reactions were incubated at 23°C for 30 min prior to measurement of parallel and perpendicular intensities (excitation: 482 nm; emission: 530 nm), and data were plotted in GraphPad Prism 10, and where appropriate, curves were fit to a 4 parameter logistic equation Y= Bottom + (Top-Bottom)/(1+(IC_50_/X)^HillSlope^)).

### In vitro pulldown

Fifty microliters of magnetic anti-FLAG M2-beads (Sigma) (washed 3x with binding buffer containing 20 mM HEPES pH 7.5, 100 mM KCl, 5 mM MgCl_2_, 1 mg/mL BSA, 1 mM TCEP pH 7.0) were incubated with various amounts of purified trimeric (FLAG-tagged) eIF2 or eIF2-P for 1h at 4°C on a rotator. Supernatans was removed and beads were washed once in binding buffer, prior to adding eIF2B to a final concentration of 2 nM and incubating once more at 4°C for 1h on a rotator. The supernatans, containing unbound protein, was partially combined with Laemmli buffer, denatured, and run on SDS-PAGE and Western Blot. Blots were probed for the various eIF2B subunits to measure and calculate the fraction bound to immobilized eIF2 (for which at least 95% always bound to the beads). Total fraction eIF2B bound was calculated for each subunit separately (β-γ-δ-ε) and then averaged and fit to the binding equation Y = Bmax*X/(K_D_ + X) using GraphPad Prism 10.

### Cryo-EM

#### Sample preparation and data collection

eIF2B decamers were assembled on ice for 15 minutes by combining eIF2B(βγδε) tetramers (WT or δL179V) with eIF2Bα2 at final concentrations of 12 µM and 9 µM, respectively. Where applicable, ligands were added after complex assembly at the following molar ratios to decamer: 15:1 ISRIB (0.5% DMSO), 15:1 mISRIB(R,R) (0.5% DMSO). Following a 30 minute incubation on ice with ligands, samples were vitrified using a Vitrobot Mark IV plunge freezer (Thermo) set to 4 °C and 95% humidity. Quantifoil R 1.2/1.3, Au 300 mesh 25 nm gold grids were glow discharged using an EasiGlow glow discharger (Pelco) for 60 s at 15 mA immediately before use. 3.5 µL of sample was applied to a grid and excess liquid was removed using dual-side blotting with a blot force of –5 for 1.5 s. The grids were immediately plunged into liquid ethane then transferred to and stored in liquid nitrogen until clipping and imaging. Datasets were collected using a Titan Grios G4 TEM (Thermo) operating at 300 keV equipped with Falcon 4i detectors and SelectrisX energy filter with a slit width of 10 eV. Images were collected at a pixel size of 0.9286 Å, with a total dose of 50 e-/Å2 across 50 frames using EPU. Full details of microscope and data collection parameters for each dataset are provided in Suppl. Table 1.

#### Cryo-EM image analysis

For all datasets, patch-based global and local motion correction and patch CTF estimation were performed within cryoSPARC live using default settings. The δL179V-ISRIB micrographs were further processed by denoising prior to template picking. Particle picking was carried out using templates generated from a published eIF2B volume (EMD-23209) particle diameter of 180 Å and minimum separation distance of 0.5 diameters. Particles were extracted with a box size of 400 pixels and Fourier cropped to 100 pixels, then subjected to 2D classification into 100 classes with a batch size per class of 400, 40 online-EM iterations and 3 full iterations. Classes resembling eIF2B were selected and the particles were subjected to ab initio reconstruction and heterogeneous refinement. Particles which yielded classes with good orientation distribution were selected, re-extracted without Fourier cropping, and refined by homogeneous or non-uniform refinement. Particles were further processed by local and global CTF refinement, and reference-based motion correction. These particles were next subjected to 3D classification and heterogenous reconstruction to generate final maps used for model building and analysis. The data processing pipeline is also outlined in Suppl. Fig. 4-6.

#### Model building, refinement and analysis

Initial models were generated from PDB 9Y3P for A-state structures and PDB 9Y3Q for I-state structures. These models were fit into experimental maps by global relaxation using ISOLDE within UCSF ChimeraX by running the simulation with simple distance restraints enabled. Each chain and residue were then manually refined to fit the experimental map. Models for ISRIB and mISRIB(R,R) were produced from their SMILES codes using AceDRG then fit into the maps with ISOLDE. A final refinement of all models was carried out against half-maps using Servalcat and the resulting statistics and MolProbity scores are reported in Supplemental table X. Experimental structures are deposited to the wwPDB with accession codes PDB: 10GE, 10GF, 10GG, 10GD; EMD-75148, 75149, 75150, 75147. All figures including cryo-EM maps and models were generated using ChimeraX.

### Generation of human cells with endogenous δL179V mutation

Editing of the *EIF2B4* gene to introduce the L179V mutation in HEK293-Flp-In-T-Rex cells was performed using CRISPR-Cas9 as described in Boone et al ^48^. Cells were seeded at 250,000 cells/well of a 12-well plate and grown for 24 h prior to transfection with a PAGE-purified, phosphorothioate-protected single-stranded oligonucleotide donor (ssODN) for homologous recombination (C016) (Renaud et al. 2016), a plasmid containing Cas9-GFP, and a plasmid encoding the guide RNA (MLM3636-C006). The 100 nt ssODN was designed to simultaneously introduce the L179V missense mutation (CTC to GTC), to remove a BamHI restriction site at S175 (GGATCC to GCTTCC), and to block re-digestion by Cas9 after recombination. Transfection was done with Xtreme Gene9 reagent according to the manufacturer’s protocol, using a 3:1 ratio of reagent (µl) to DNA (µg). Reagent-only and pCas9-GFP controls were included. Two days post transfection, cells were trypsinized, washed twice in ice-cold filter-sterilized FACS buffer (25 mM HEPES pH 7.0, 2 mM EDTA, 0.5% v/v fetal bovine serum, in 1x PBS), and resuspended in FACS buffer with 400 ng/ml 7-AAD viability dye (Invitrogen) at around 1 million cells/ml in filter-capped FACS tubes. Single GFP^+^, 7-AAD^−^ cells were sorted into recovery medium (a 1:1 mix of conditioned medium, and fresh medium with 20% fetal bovine serum, 2 mM L-Glutamine, 1 mM sodium pyruvate, and 1x non-essential amino acids) in single wells of 96-well plates using the Sony SH800 cell sorter. The survival rate was around 4% after 2-3 weeks. Surviving clones were expanded and first screened for correct editing by PCR and BamHI restriction digest. For this, genomic DNA was isolated using the PureLink Genomic DNA mini kit (Invitrogen), and a 437 bp fragment of the *EIF2B4* gene was amplified by PCR using 300 nM forward and reverse primers (C002_F and C002_R), 300 µM dNTPs, 1x HF buffer, 100 ng genomic DNA / 100 µl reaction and 2 U/100 µl reaction of KAPA HiFi polymerase for 3 min at 95°C; and 30 cycles of 98°C for 20 s, 68.9°C for 15 s, 72°C for 15 s, prior to cooling at 4°C. PCR reactions were purified using NucleoSpin Gel and PCR cleanup kit (Macherey Nagel), and HighPrep PCR Cleanup beads (MagBio Genomics) using the manufacturer’s instructions. Cleaned up products were digested using BamHI restriction enzyme (NEB) in 1x CutSmart buffer and run on a 1.5% agarose gel with 1x SYBR Safe (Invitrogen) and 100 bp DNA ladder (Promega). Clones that lost the BamHI restriction site were then deep sequenced to confirm correct editing and zygosity. For this, the *EIF2B4* gene was amplified by PCR using 300 nM forward and reverse primers (C035_F and C035_R), 300 µM dNTPs, 1x HF buffer, 100 ng genomic DNA / 100 µl reaction and 2 U/100 µl reaction of KAPA HiFi polymerase for 3 min at 95°C; and 30 cycles of 98°C for 20 s, 63°C for 15 s, 72°C for 15 s, prior to cooling at 4°C. The 200 bp product was purified from a 1.5% agarose gel with 1x SYBR Safe using NucleoSpin Gel and PCR cleanup kit (Macherey Nagel), and HighPrep PCR Cleanup beads (MagBio Genomics) using the manufacturer’s instructions. A subsequent second PCR added the Illumina P5/P7 sequences and barcode for deep sequencing. For this, we used 15 ng purified PCR product per 100 µl reaction, 300 nM forward and reverse primer (C037_F_bcx, and C037_R), and 1x KAPA HiFi HotStart mix, for 3 min at 95°C, and 8 cycles of 20 s at 98°C, 15 s at 66.3°C, and 15 s at 72°C prior to cooling on ice. PCR reactions were purified using HighPrep beads (MagBio Genomics), and amplicon quality and size distribution was checked by chip electrophoresis (BioAnalyzer High Sensitivity kit, Agilent). Samples were then sequenced on an Illumina MiSeq (150 bp paired-end), and results were analyzed with CRISPResso2 (Pinello et al. 2016). All cell lines were negative for mycoplasma contamination.

### Generation of ATF4-mVenus reporter cell lines

Vesicular stomatitis virus-G pseudotyped lentivirus containing pMB033 were prepared by transfecting 293T cells with 2.5 ug pMB033, 2 ug pMD2.G, and 2 ug psPAX2 plasmids in OptiMEM using Lipofectamine 2000, according to the manufacturer’s instructions. Cells were incubated with the transfection mix for 6h at 37C prior to replacing the medium with DMEM. Twenty-four hours after transfection, medium was replaced with collection medium (DMEM containing 4.5 g/L glucose supplemented with 10% FBS, 15 mM HEPES, L-glutamine, and sodium pyruvate). Forty-eight hours after transfection (or 24h after medium replacement), the medium containing the viral supernatant was spun down to remove dead cells, filtered through a 0.45 μm (low protein binding) filter unit (EMD Millipore) to remove debris, and concentrated with Lenti-X concentrator solution (1 volume concentrator per 3 volumes viral medium). After overnight incubation at 4C, viral particles were centrifuged for 45 min at 4C and 1500g, resuspended in ice-cold PBS, aliquoted and frozen to –80C for future use. For spinfection, target WT or L179V cells were seeded at 400,000 cells per well of a 6-well plate and grown for 24h. Medium was exchanged for medium with 8 μg/ml polybrene, virus was added, and the plate was centrifuged for 2h at 700g at 23C. After 24h of incubation at 37C and 5% CO_2_, virus-infected supernatans was removed, and cells were gently washed with warm PBS twice before returning to fresh medium. Cells were allowed to recover for another 48h, and split if reached 70% confluency before that. Roughly 100,000 TagBFP2-positive cells were then sorted on a Sony MA900 cytometer to select for those that integrated the overexpression plasmid, and were subsequently expanded and frozen as polyclonal reporter cell lines.

### Growth Curves

Cells were seeded in a 6-well plate at 200,000 cells/well +/− 200 nM (m)ISRIB with final 0.1% DMSO, and grown at 37°C and 5% CO_2_. At 70% confluency (2 days later), cells were trypsinized, counted, and 1/6 were transferred to a new well of a 6-well plate. This was repeated for 10-12 days. Final cell counts were corrected for the cumulative dilution factor.

### IncuCyte live-cell imaging

For all IncuCyte experiments with HEK293 cells, black-walled 96-well plates with glass bottom (Cellvis P96-1.5H-N) were coated with 100 ug/ml Poly-D Lysine (PDL, Fisher Scientific) for 1h at 37°C, washed 3x with PBS, and thoroughly air dried prior to RT storage. Cells were seeded in medium at 5,000 cells per well (100 ul per well) and allowed to recover for 24h at 37°C. A pre-treatment baseline measurement was done in the Incucyte 30 mins after plate introduction, after which 50% of the medium was gently replaced with a 2x mix containing indicated drugs (including Cytotox NIR Dye at 1.2 µM), with a final DMSO of 0.1%. Cells were immediately placed back in the IncuCyte S3 Live-Cell Analysis System (Sartorius) and allowed to recover for 30 mins prior to the first measurement. Cells were imaged every 2h for 5 days with a 10x objective, with an adherent cell-by-cell scan type using 3 images per well per timepoint, using the phase contrast, green, and NIR fluorescence channels. Image analysis was done in the IncuCyte Analysis Software. Cell confluence measurement was performed using the AI confluence algorithm setting a min. of 300 µm^2^ per cell. For fluorescent and cell death reporters, we used a cell-by-cell based analysis, accounting for a 2% contribution of the NIR signal to the green signal, and setting a min. cell size of 300 µm^2^. For counting the fraction of dead cells, 2 cell classes were defined (alive/dead) with 0.2 NIRCU value as cut-off.

### Western Blotting

*Cultured cells*. Western blotting was done as in Boone et al^48^. Cells were seeded at 400,000 cells/well of a 6-well plate and grown at 37°C and 5% CO_2_ for 24 h. For drug treatment, final DMSO concentration was 0.1% across all conditions. For the protein synthesis assay, cells were treated with drugs as indicated for 1h, and a puromycin pulse (10 µg/ml) was given during the last 10 min of incubation. Plates were put on ice, cells were washed once with ice-cold phosphate-buffered saline (PBS), and then lysed in 150 μl ice-cold lysis buffer (50 mM Tris-HCl pH 7.4, 150 mM NaCl, 1 mM EDTA, 1% v/v Triton X-100, 10% v/v glycerol, 1x cOmplete EDTA-free protease inhibitor cocktail [Roche], and 1x PhosSTOP [Roche]). Cells were scraped off, collected in an eppendorf tube, and put on a rotator for 30 min at 4°C. Debris was pelleted at 12,000 g for 20 min at 4°C, and supernatant was removed to a new tube on ice. Protein concentration was measured using the bicinchonic acid (BCA) assay. Within an experiment, total protein concentration was normalized to the least concentrated sample (typically all values were within ∼10%). A 5x Laemmli loading buffer (250 mM Tris-HCl pH 6.8, 30% glycerol, 0.25% bromophenol blue, 10% SDS, 5% β-mercaptoethanol) was added to each sample to 1x, and samples were denatured at 95°C for 12 min, then cooled on ice. Wells of AnyKd Mini-PROTEAN TGX precast protein gels (AnyKD, Bio-Rad) were loaded with equal amounts of total protein (10-15 µg), in between Precision Plus Dual Color protein ladder (BioRad). After electrophoresis, proteins were transferred onto a nitrocellulose membrane at 4°C, and then blocked for 2h at room temperature in PBS with 0.1% Tween (PBS-T) + 3% milk (blocking buffer) while rocking. Primary antibody staining was performed with gentle agitation at 4°C overnight. After washing four times in blocking buffer, secondary antibody staining was performed for 1h at room temperature using anti-rabbit HRP or anti-mouse HRP secondary antibodies (Promega, 1:10,000) in blocking buffer. Membranes were washed 3x in blocking buffer and then 1x in PBS-T without milk. Chemiluminescent detection was performed using SuperSignal West Dura or Femto HRP substrate (Thermo Fisher Scientific), and membranes were imaged on a LI-COR Odyssey gel imager for 0.5–10 min depending on band intensity.

For the phospho-retention blots, equal amounts of total protein lysates (10-15 µg) were loaded on 12.5% Supersep Phos-tag gels (Wako Chemicals) in between Wide-view III protein ladder (Wako Chemicals). After electrophoresis, the gel was washed 3x in transfer buffer with 10 mM EDTA prior to transfer onto nitrocellulose. Blocking, antibody staining and detection was performed as described above.

*Mouse brain tissues*. Mouse hippocampus and cortex from each hemisphere were dissected on ice and stored at –80°C. For lysis, tissues were manually homogenized in freshly made cold homogenizing buffer (200 mM HEPES, 50 mM NaCl, 10% Glycerol, 1% Triton X-100, 1 mM EDTA, 50 mM NaF, 2 mM Na_3_VO_4_, 25 mM β-glycerophosphate, and 1x cOmplete EDTA-free protease inhibitor cocktail [Roche) with a sterile pestle, and passed through a p200 pipette tip. Samples were then put on a rotator at 4°C for 15 min, and centrifuged at 12,000 rpm for 20 min at 4°C. The supernatant was transferred to a fresh tube. Protein concentration of 10x diluted samples was measured using the bicinchonic acid (BCA) assay, and normalized to the sample with lowest concentration. Western Blotting was further done as described for cultured cells, but loading 20-25 µg of protein per sample, and using goat or donkey anti-rabbit or anti-mouse IRDye 800CW secondary antibodies (LICORBio, 1:10,000) in blocking buffer for secondary staining followed by fluorescent detection. Quantification was done with the EmpiriaStudio software.

### Mouse liver microsome assay

Mouse liver microsome assays were performed by Quintara Biosciences (Hayward, CA). Reactions contained a final concentration of 1 μM ISRIB/mISRIB/Verapamil (positive control), 0.5 mg/mL liver microsome protein, and 1 mM NADPH in 100 mM potassium phosphate, pH 7.4 buffer with 3 mM MgCl_2_ in a final 25 μL volume. At each of the time points (0, 15, 30, and 60 minutes), 150 μL of quench solution (acetonitrile with 0.1% formic acid) with internal standard (bucetin) was transferred to each well. Plates were centrifuged at 4°C for 15 minutes at 4000 rpm, and supernatant was analyzed by LC-MS/MS analysis. Metabolism was calculated as the fraction of test compound remaining compared to t_0_. Initial rates were calculated for the compound concentration and used to determine t_1/2_ values and the intrinsic clearance (Clint).

### Mouse husbandry

To generate the constitutive mice with the *Eif2b4-L180V* mutation at the endogenous locus, we crossed CMV-Cre mice (Jackson Laboratory strain #006054, rederived at Altos Labs, C57Bl/6J background) with mice with a floxed WT minigene cassette inserted into exon 3 and the L180V mutation in exon 6 at the endogenous *Eif2b4* locus (generated by Ingenious Labs, mixed C57Bl6/N and C57Bl/g-FLP background). Mice with germline transmission of the (now constitutive, and cassette-less) *Eif2b4-L180V* allele were further backcrossed with wild-type C57Bl/6J mice for several generations. Mice were genotyped by Transnetyx (Cordova, TN) from tail biopsies using real-time PCR with specific probes designed for each allele.

All *in vivo* experiments were conducted on 3-4 months old mice. Mice were housed on a 12-hr light/dark cycle (lights on at 7:00 am) in a temperature– and humidity-controlled environment, with all experiments and procedures performed during the light phase of the cycle. Animals had *ad libitum* access to standard chow and water. Animal studies were approved by the Charles River Laboratories’ and Altos Labs’ Institutional Animal Care and Use Committee in accordance with guidelines established by the National Institutes of Health.

### Mouse drug administration

ISRIB or mISRIB solutions for mouse ip injections were prepared fresh daily, at a final concentration of 0.1 mg/ml (∼222 μM) and injected at 0.25 mg/kg. To this end, 100 ul aliquots of 2 mg/mL (∼4.4 mM) ISRIB or mISRIB solutions in DMSO (stored at –80°C) were freshly thawed at 37°C, and mixed well with 40 μl Tween 80 (Sigma Aldrich, P8074). The solution was gently heated to 37°C and vortexed until the solution was clear. Next, 400 μl of polyethylene glycol 400 (PEG400) (MedChemExpress HY-Y0873A) solution was added, and again gently heated to 37°C and vortexed until the solution was clear. Finally, 1,460 μl of 5% dextrose was added and once more heated to 37°C and vortexed until the solution was clear. The solution was kept at room temperature throughout the experiment. The vehicle solution was prepared in the same way, replacing the ISRIB/mISRIB with 100% DMSO.

### Mouse tissue collection

For pharmacokinetics studies, mice were anesthetized with avertin (2,2,2-Tribromoethanol, Sigma T48402). Once animals were completely anesthetized, blood was extracted by cardiac puncture and animals were perfused with 0.9% NaCl until the livers were clear (∼10 mL). Whole brain was rapidly removed, regions of interest were dissected on ice, snap frozen in liquid nitrogen, and stored at −80°C until processing. For plasma processing, ∼70 µl blood was collected in EDTA-coated tubes, mixed well, and kept on ice for 15 min. Blood was then centrifuged at 1500g for 15 min at 4°C and 20 µl of the top layer was recovered as plasma, flash frozen in liquid nitrogen, and stored at or below – 80°C until processing.

For regular tissue collection, mice were euthanized with CO_2_ and cervical dislocation, and brain tissues were processed as above.

### *In vivo* pharmacokinetics

*Sample Preparation*. Plasma samples were extracted using LC-MS grade methanol (MeOH) at a 10:1 (MeOH:plasma) ratio. Following extraction, samples were centrifuged, and the resulting supernatant was transferred to a glass injection plate. A 1 µL aliquot of each sample was injected onto the mass spectrometer. Tissue samples were extracted using LC-MS grade MeOH at a ratio of 9 µL MeOH per 1 mg tissue. The extraction solvent was added to the tissue, homogenized using a Geno/Grinder, and centrifuged. The supernatant was then transferred to a glass injection plate, and 1 µL was injected onto the mass spectrometer.

*LC-MS/MS Analysis*. Samples were analyzed on a SCIEX Triple Quad 7500 operated in negative ion mode. The monitored transitions were mISRIB 479.1 → 154 and ISRIB 450.9 → 186. Chromatographic separation was performed using a Thermo Vanquish Duo UHPLC equipped with a Waters BEH C18 2.1×100mm 1.7µm column maintained at 50 °C. Mobile phase A consisted of 0.15% formic acid in water, and mobile phase B consisted of 0.1% formic acid in acetonitrile (ACN). The gradient program was as follows: 0–0.5 min, 25% B at 0.40 mL/min; 0.5–4.5 min, 100% B at 0.40 mL/min; 4.5–5.5 min, 100% B at 0.60 mL/min; 5.5–6 min, 25% B at 0.60 mL/min; and 6–6.5 min, 25% B at 0.40 mL/min. The total run time was 6.5 minutes.

*Calibration Curve*. Calibration curves were generated by spiking mISRIB and ISRIB standards (prepared in MeOH) into extracted pooled plasma and brain samples. For plasma, 5 µL of standard was diluted into 45 µL of pooled extracted plasma. For brain tissue, standards were prepared by serial dilution in MeOH and then spiked into extracted matrix. Final concentrations of mISRIB and ISRIB in the calibration samples were 0, 10, 20, 50, 100, 200, 500, and 1000 ng/mL. Each calibration sample was analyzed by injecting 1 µL onto the mass spectrometer.

*Quantification*. Peak areas were determined using SCIEX OS software. The baseline peak area measured from the 0 ng/mL calibration sample was subtracted from all samples. Known standard concentrations and their measured peak areas were used to construct a linear calibration curve (line of best fit). The concentrations (ng/mL) of mISRIB and ISRIB in unknown samples were calculated using this calibration curve.

### Transcriptomics of cortical tissues

Frozen mouse hemicortices were manually homogenized in 700 ul TriZOL with RNAse-free pestles, passed 10x through an RNAse-free needle, and debris was spun down for 5 mins at 3200x g. Total RNA was then extracted from the supernatans using the DirectZOL RNA mini kit (Zymo) with DNase digestion according to the manufacturer’s instructions. Concentration was checked with the Qubit RNA Broad Range assay kit (Thermo Scientific), and total RNA quality was confirmed (RIN >= 8) using the TapeStation RNA ScreenTape assay (Agilent) according to the manufacturer’s instructions. PolyA-sequencing libraries were prepared using the Watchmaker Genomics mRNA library prep, quantified using qPCR and the KAPA Library Quantification kit (Roche), and samples were sequenced on an Illumina NextSeq 500 at 30 million reads per sample. Reads were demultiplexed, analyzed with FastQC v0.11.9, trimmed with TrimGalore 0.6.7 and Cutadapt 3.4, and aligned to the Ensembl *Mus musculus* GRCm39 genome with Salmon v1.28.0. Length-scaled read counts were used as input for differential gene expression data analysis with DESeq2 in R (v4.5.2) with bioconductor (v3.22). Data was deposited in GEO with accession number XXX.

### Electrophysiology

Electrophysiological recordings were performed as previously described^23^. Briefly, field recordings were performed from CA1 horizontal hippocampal slices (320 µm thick), which were cut from the brain of adult mice (3-4 months old) with a vibratome (Leica VT 1200S, Leica Microsystems, Buffalo Grove, IL) at 4°C in artificial cerebrospinal fluid solution (ACSF; 95% O_2_ and 5% CO_2_) containing 124 mM NaCl, 2.0 mM KCl, 1.3 mM MgSO_4_, 2.5 mM CaCl_2_, 1.2 mM KH_2_PO_4_, 25 mM NaHCO_3_, and 10 mM glucose (2-3 ml/min). Slices were incubated for at least 60 min prior to recording in an interface chamber and continuously perfused with artificial cerebrospinal fluid (ACSF) at 28 – 29°C at a flow rate of 2-3 ml/min. The recording electrodes were placed in the stratum radiatum. Field excitatory postsynaptic potentials (fEPSPs) were recorded with ACSF-filled micropipettes and were elicited by bipolar stimulating electrodes placed in the CA1 stratum radiatum to excite Schaffer collateral and commissural fibers. The intensity of the 0.1-ms pulses was adjusted to evoke 30 – 35% of maximal response. A stable baseline of responses at 0.033 Hz was established for at least 30 min. L-LTP was induced by applying a single high-frequency stimulus (100 Hz, 1 s). Slices were treated with vehicle (0.05% DMSO), ISRIB (200 nM), or mISRIB (400 nM) for at least 30 min before high-frequency stimulation and were maintained in the same solution throughout the recording. Concentrations were empirically determined based on their capacity to enable L-LTP induction: vehicle (0.05% DMSO) did not elicit L-LTP, while 200 nM ISRIB and 400 nM mISRIB both enabled robust L-LTP. The ISRIB concentration used exceeds the 50 nM concentration previously shown to be subthreshold for L-LTP induction^17^. For each mouse, two slices were used per recording and counted as independent measurements. For WT + ISRIB samples, one recording failed and was discarded. Statistical comparisons were calculated on the average of the last 5 timepoints of the measurements (2h after HFS) for each replicate.

### Behavioral assays

For all behavioral experiments, the experimenter was blinded to genotype and treatment, and i.p. injections were performed by a different experimenter. Mice were handled one by one for 5 min per day for 3 days prior to the experiment(s). When multiple behavioral tests were to be run on the same cohort, they were performed in the order of least to most stressful. Minimal sample sizes were estimated based on previously published work and our preliminary investigations. Post-hoc power analysis demonstrated that the sample sizes used were sufficient to detect an effect size of 1.1 at 80% power and alpha = 0.05 (two-sided t-test).

#### Open Field (OF)

Mice were placed a sound-attenuating isolation cubicle (w-d-h: 25.5 cm x 25.5 cm x 36 cm, Mouse Cage XL from Ugo Basile) for 20 min per day for 3 days, with running fan and dim lights. To control for odor cues, the cubicle was thoroughly cleaned with ethanol, dried, and ventilated between mice. Mouse movement was recorded from cameras positioned above the training chamber, and distance traveled was calculated using the Noldus Ethovision XT version 17 automated tracking software. Locomoter activity was calculated as the distance traveled during habituation, averaged over 3 days.

#### Contextual fear conditioning (CFC)

Mice were first habituated to a sound-attenuating isolation cubicle (w-d-h: 25.5 cm x 25.5 cm x 36 cm, Mouse Cage XL from Ugo Basile) for 20 min per day for 3 days, with running fan and dim lights. The training protocol varied depending on the goal of the experiment: a weak training protocol was followed to enable detection of enhanced freezing (ie long-term fear memory enhancement) compared to baseline, and a more conventional strong training protocol to better detect reduced freezing (ie long-term fear memory impairment). On the training day, mice were placed in the cubicle for 2 min (naïve) and then received either one ‘weak’ foot shock (0.35 mA, 1s for the weak protocol) or two ‘strong’ foot shocks (0.75 mA, 2s, 90s apart for the strong protocol), after which the mice remained in the chamber until the 5 min mark before being returned to their home cages. Twenty-four hours later, mice were re-exposed to the same context (the chamber) for 5 min. Mouse movement was recorded from cameras positioned above the training chamber, and freezing (immobility with the exception of respiration) was calculated using the Noldus Ethovision XT version 17 automated tracking software. The fear condition training was always the last behavioral test performed.

## Acknowledgments

We thank Calico for the initial generous gift of purified eIF2 heterotrimer to UCSF; Matias Hartman from the Denzel lab (Max Planck Institute for the Biology of Ageing) for the mouse ES cells with homozygous L180V mutation, Lucas Reineke for sharing CReP mutant mice, and the Altos BAI Genomics Hub for RNASeq library prep and sequencing.

## Author contributions

M.B. conceptualized the project, performed experiments, analyzed the data, and wrote the manuscript. U.D., A.D., P-J.Z., T.C., M.Y., K.P., R. M., I.B., J.W., H.Z., D.J.L., C.P.A., T.G.L., H.Z., M.K., P.E., M.S., R.L. performed additional experiments or provided experimental support. P.W. and A.F. conceptualized and supervised the project. M.C.M. and A.R. provided additional supervision and advice. All authors read, revised and approved the final version of the manuscript.

## Data and materials availability

Cryo-EM density maps and associated models will be publicly available through the EMDB with accession 75148, 75149, 75150, 75147 and PDB with accession codes 10GE, 10GF, 10GG, 10GD respectively. RNASeq data is available on GEO with accession code XXX. All other raw data are available upon request through data transfer agreements.

