## Supplemental Figures for "Stereoselective methyl-swapping demonstrates target specificity of cognitive enhancer"

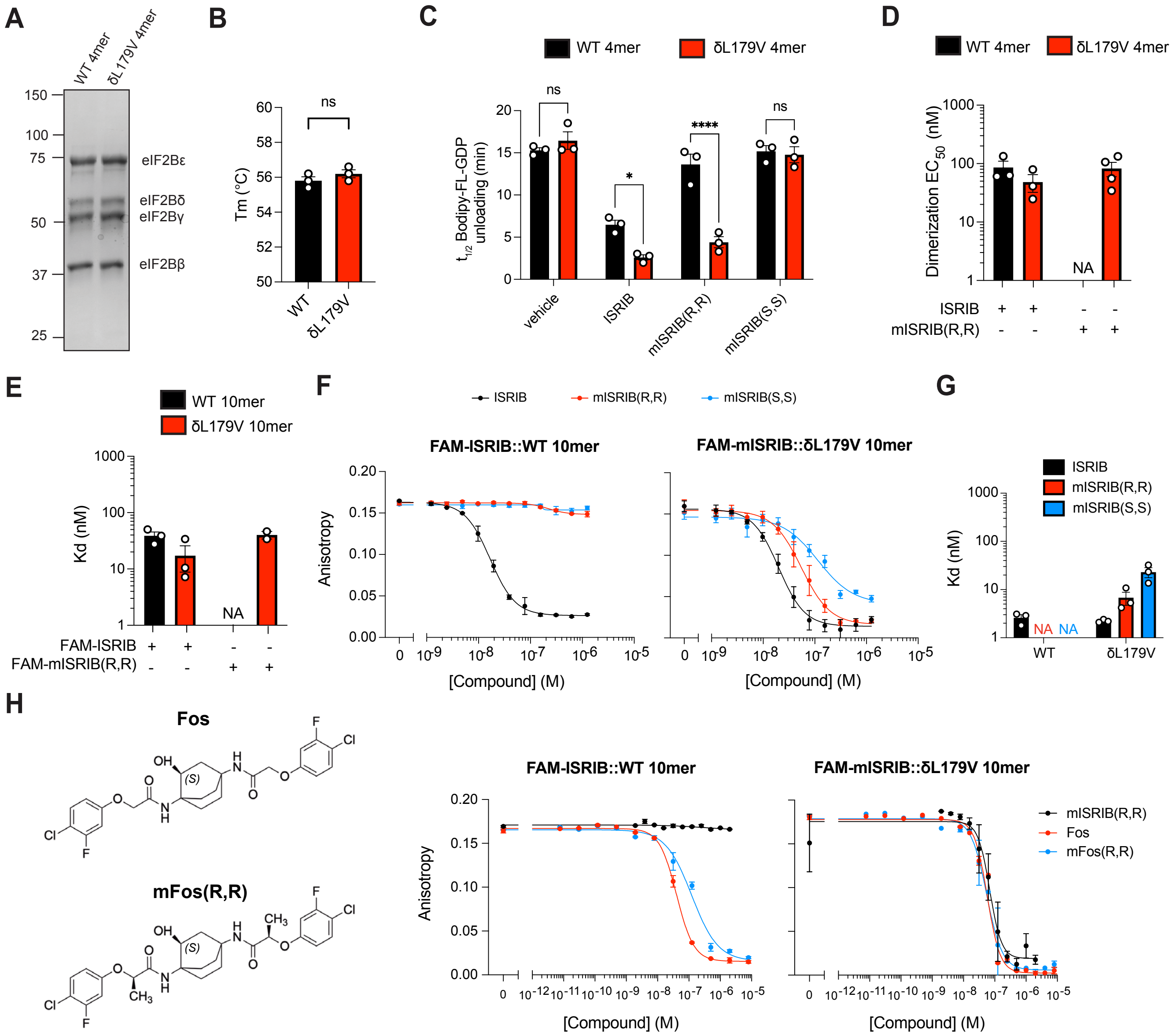
A
WT 4mer
δL179V 4mer
150
100
75
50
37
25
eIF2Bε
eIF2Bδ
eIF2Bγ
eIF2Bβ
B
ns
Tm (°C)
WT
δL179V
C
WT 4mer
δL179V 4mer
t1/2 Bodipy-FL-GDP unloading (min)
ns
*
****
vehicle
ISRIB
mISRIB(R,R)
mISRIB(S,S)
D
Dimerization EC50 (nM)
NA
ISRIB
mISRIB(R,R)
E
WT 10mer
δL179V 10mer
Kd (nM)
NA
FAM-ISRIB
FAM-mISRIB(R,R)
F
ISRIB
mISRIB(R,R)
mISRIB(S,S)
FAM-ISRIB::WT 10mer
FAM-mISRIB::δL179V 10mer
Anisotropy
[Compound] (M)
G
ISRIB
mISRIB(R,R)
mISRIB(S,S)
Kd (nM)
NA
NA
WT
δL179V
H
Fos
mFos(R,R)
FAM-ISRIB::WT 10mer
FAM-mISRIB::δL179V 10mer
Anisotropy
[Compound] (M)
mISRIB(R,R)
Fos
mFos(R,R)

Supp. Fig. 1

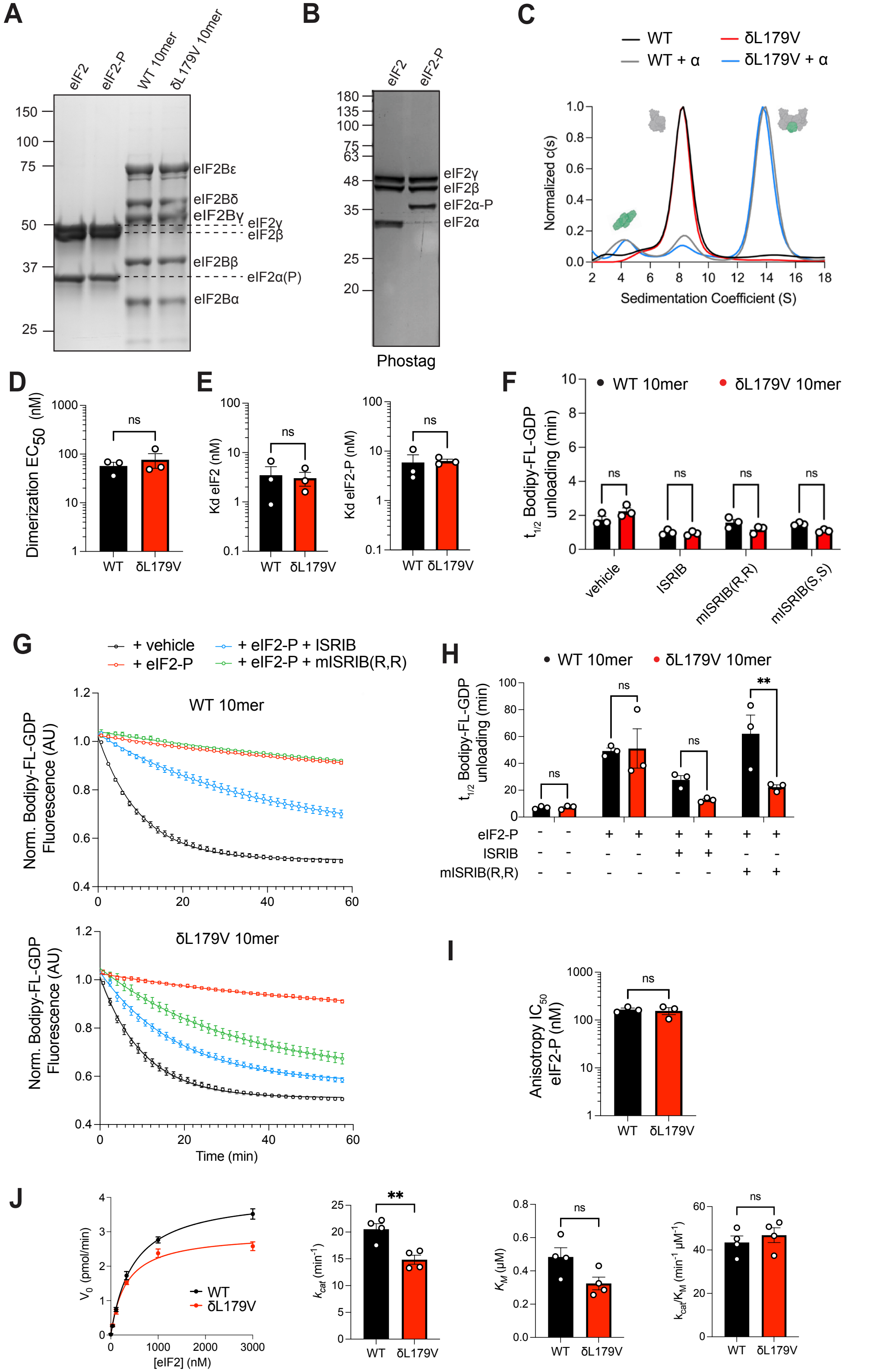
A
eIF2
eIF2-P
WT 10mer
δL179V 10mer
150
100
75
50
37
25
eIF2Bε
eIF2Bδ
eIF2Bγ
eIF2γ
eIF2β
eIF2Bβ
eIF2α(P)
eIF2Bα
B
eIF2
eIF2-P
180
135
100
75
63
48
35
25
20
eIF2γ
eIF2β
eIF2α-P
eIF2α
Phostag
C
WT
WT + α
δL179V
δL179V + α
Normalized c(s)
Sedimentation Coefficient (S)
D
Dimerization EC50 (nM)
ns
WT
δL179V
E
Kd eIF2 (nM)
Kd eIF2-P (nM)
ns
WT
δL179V
F
WT 10mer
δL179V 10mer
t1/2 Bodipy-FL-GDP unloading (min)
ns
vehicle
ISRIB
mISRIB(R,R)
mISRIB(S,S)
G
+ vehicle
+ eIF2-P
+ eIF2-P + ISRIB
+ eIF2-P + mISRIB(R,R)
WT 10mer
δL179V 10mer
Norm. Bodipy-FL-GDP Fluorescence (AU)
Time (min)
H
WT 10mer
δL179V 10mer
t1/2 Bodipy-FL-GDP unloading (min)
ns
**
eIF2-P
ISRIB
mISRIB(R,R)
I
Anisotropy IC50 eIF2-P (nM)
ns
WT
δL179V
J
V0 (pmol/min)
[eIF2] (nM)
WT
δL179V
kcat (min-1)
**
KM (μM)
kcat/KM (min-1 μM-1)
ns

**Suppl. Fig. 2**

A

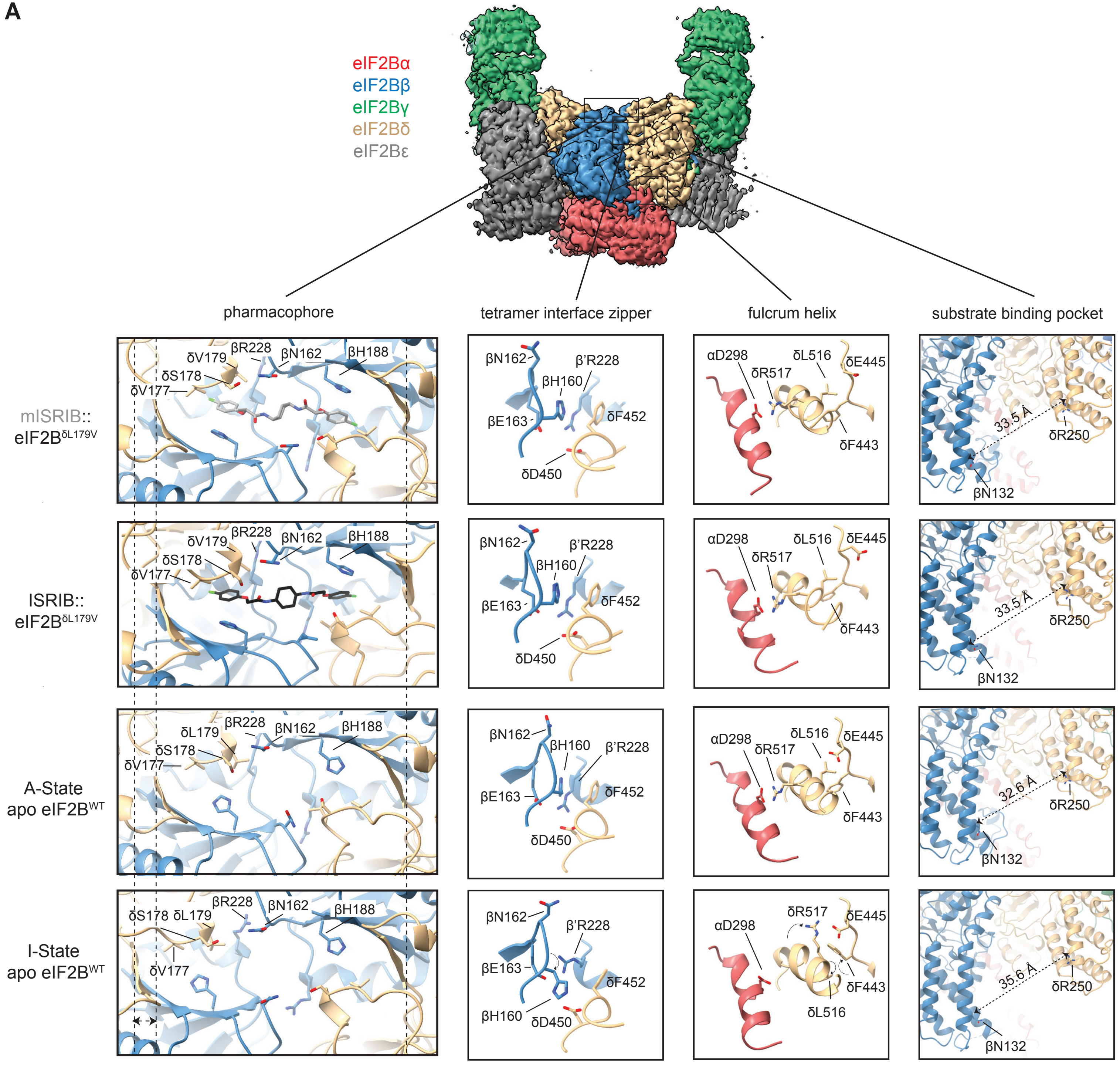

eIF2Bα
eIF2Bβ
eIF2Bγ
eIF2Bδ
eIF2Bε
pharmacophore
tetramer interface zipper
fulcrum helix
substrate binding pocket
mISRIB::
eIF2BδL179V
ISRIB::
eIF2BδL179V
A-State
apo eIF2BWT
I-State
apo eIF2BWT
δV177
δS178
δV179
δL179
βR228
βN162
βH188
β'R228
βH160
βE163
δF452
δD450
αD298
δR517
δL516
δE445
δF443
33.5 Å
32.6 Å
35.6 Å
δR250
βN132


**Suppl. Fig. 3**

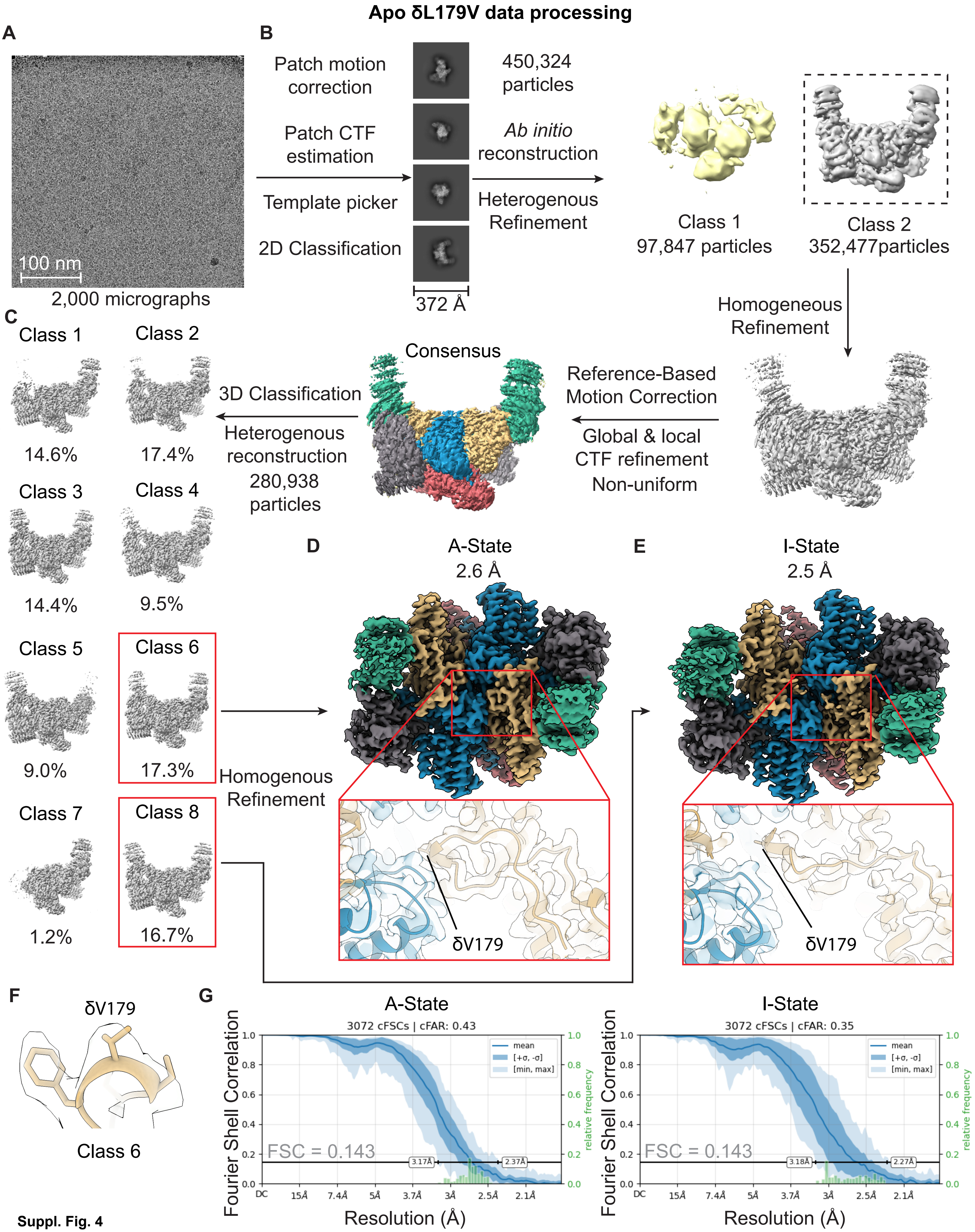
Apo δL179V data processing
A
B
Patch motion correction
Patch CTF estimation
Template picker
2D Classification
450,324 particles
Ab initio reconstruction
Heterogenous Refinement
100 nm
2,000 micrographs
372 Å
Class 1
97,847 particles
Class 2
352,477particles
Homogeneous Refinement
C
Class 1
Class 2
14.6%
17.4%
Class 3
Class 4
14.4%
9.5%
Class 5
Class 6
9.0%
17.3%
Class 7
Class 8
1.2%
16.7%
Consensus
3D Classification
Heterogenous reconstruction
280,938 particles
Reference-Based Motion Correction
Global & local CTF refinement
Non-uniform
Homogenous Refinement
D
A-State
2.6 Å
E
I-State
2.5 Å
δV179
δV179
F
δV179
Class 6
G
A-State
3072 cFSCs | cFAR: 0.43
I-State
3072 cFSCs | cFAR: 0.35
Fourier Shell Correlation
Resolution (Å)
FSC = 0.143
relative frequency
mean
[+σ, -σ]
[min, max]
3.17Å
2.37Å
3.18Å
2.27Å
DC
15Å
7.4Å
5Å
3.7Å
3Å
2.5Å
2.1Å

Suppl. Fig. 4

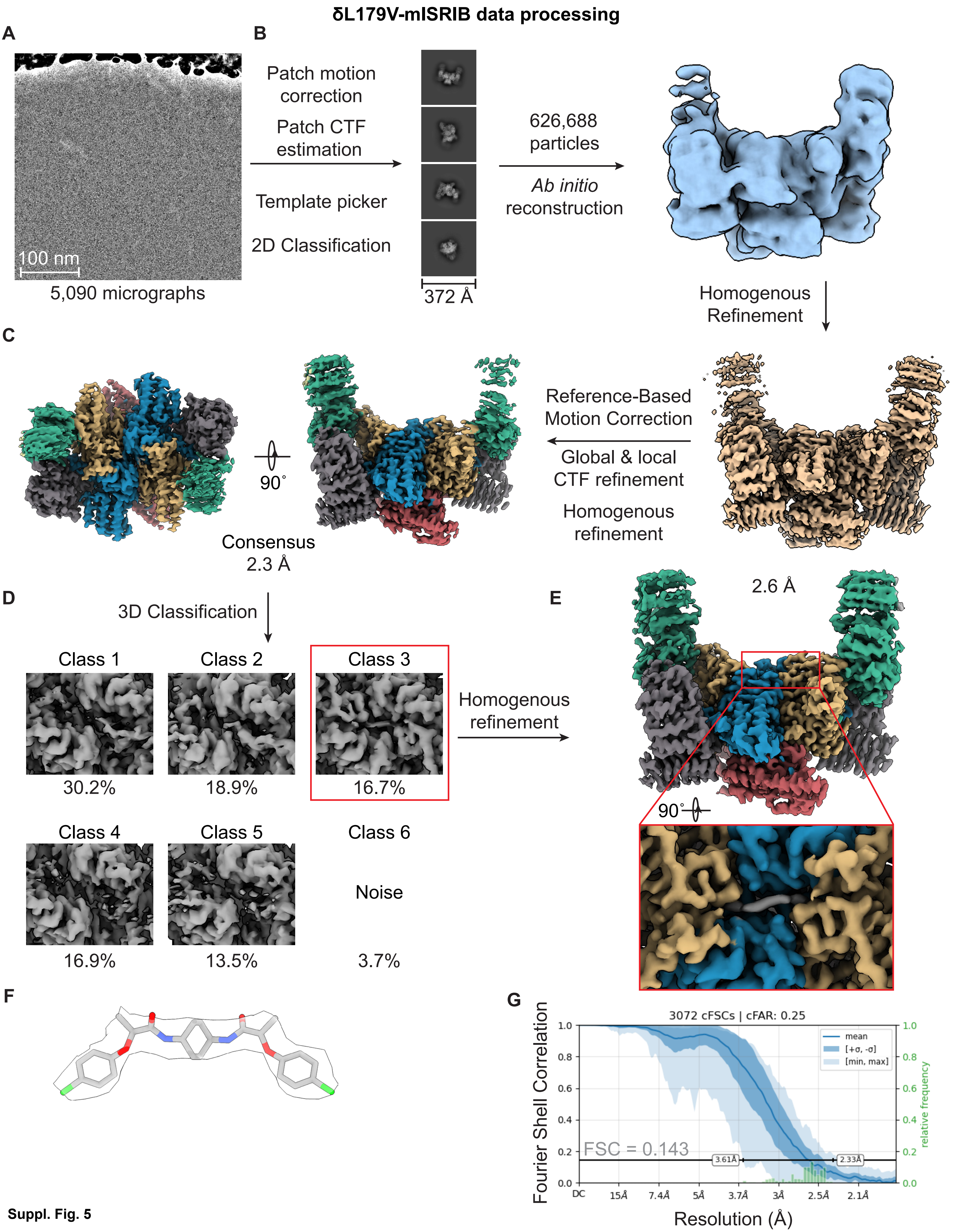

δL179V-mISRIB data processing
A
100 nm
5,090 micrographs
B
Patch motion correction
Patch CTF estimation
Template picker
2D Classification
372 Å
626,688 particles
Ab initio reconstruction
Homogenous Refinement
C
Reference-Based Motion Correction
Global & local CTF refinement
Homogenous refinement
90˚
Consensus 2.3 Å
D
3D Classification
Class 1
30.2%
Class 2
18.9%
Class 3
16.7%
Class 4
16.9%
Class 5
13.5%
Class 6
Noise
3.7%
Homogenous refinement
E
2.6 Å
90˚
F
G
3072 cFSCs | cFAR: 0.25
mean
[+σ, -σ]
[min, max]
FSC = 0.143
3.61Å
2.33Å
Fourier Shell Correlation
relative frequency
Resolution (Å)
DC
15Å
7.4Å
5Å
3.7Å
3Å
2.5Å
2.1Å


Suppl. Fig. 5

**δL179V-ISRIB data processing**

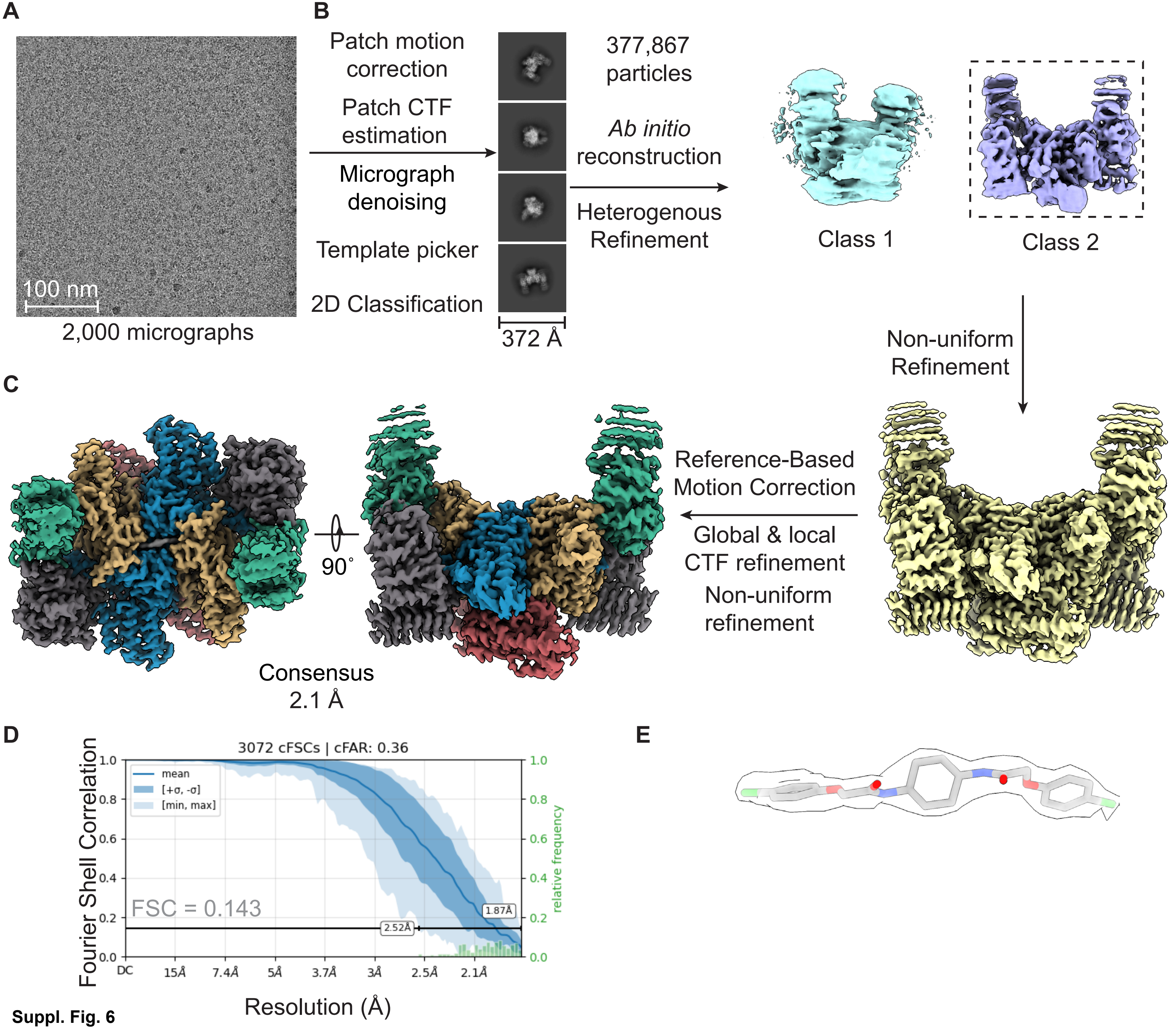


**Suppl. Fig. 6**

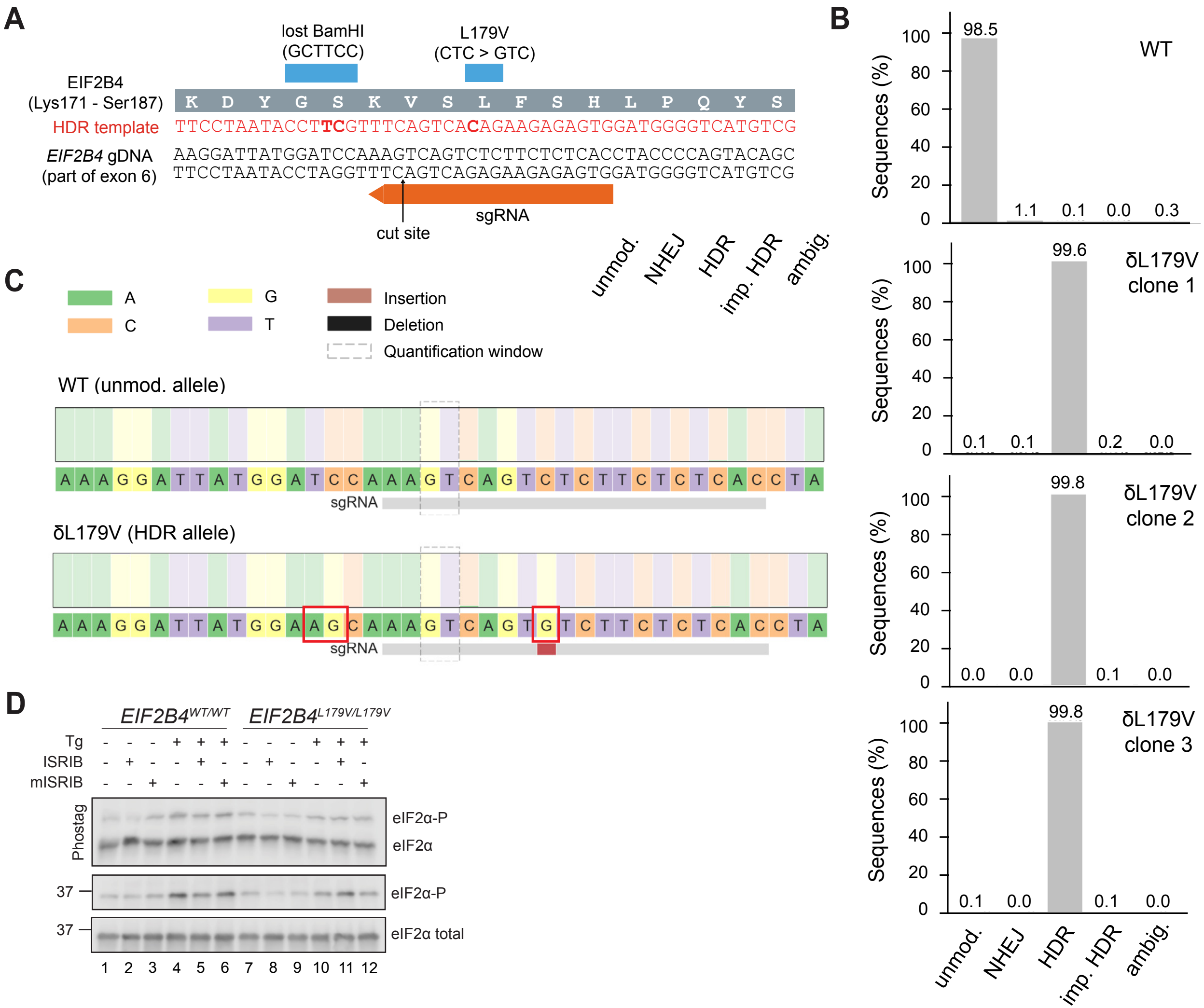
A
lost BamHI (GCTTCC)
L179V (CTC > GTC)
EIF2B4 (Lys171 - Ser187)
K D Y G S K V S L F S H L P Q Y S
HDR template
TTCCTAATACCTTCGTTTCAGTCACAGAAGAGAGTGGATGGGGTCATGTCG
EIF2B4 gDNA (part of exon 6)
AAGGATTATGGATCCAAAGTCAGTCTCTTCTCTCACCTACCCCAGTACAGC
TTCCTAATACCTAGGTTTCAGTCAGAGAAGAGAGTGGATGGGGTCATGTCG
sgRNA
cut site
B
Sequences (%)
WT
98.5
1.1
0.1
0.0
0.3
δL179V clone 1
0.1
0.1
99.6
0.2
0.0
δL179V clone 2
0.0
0.0
99.8
0.1
0.0
δL179V clone 3
0.1
0.0
99.8
0.1
0.0
unmod.
NHEJ
HDR
imp. HDR
ambig.
C
A
C
G
T
Insertion
Deletion
Quantification window
WT (unmod. allele)
A A A G G A T T A T G G A T C C A A A G T C A G T C T C T T C T C T C A C C T A
sgRNA
δL179V (HDR allele)
A A A G G A T T A T G G A A G C A A A G T C A G T G T C T T C T C T C A C C T A
sgRNA
D
EIF2B4 WT/WT
EIF2B4 L179V/L179V
Tg
ISRIB
mISRIB
Phostag
eIF2α-P
eIF2α
37
eIF2α-P
37
eIF2α total
1 2 3 4 5 6 7 8 9 10 11 12

**Suppl. Fig. 7**

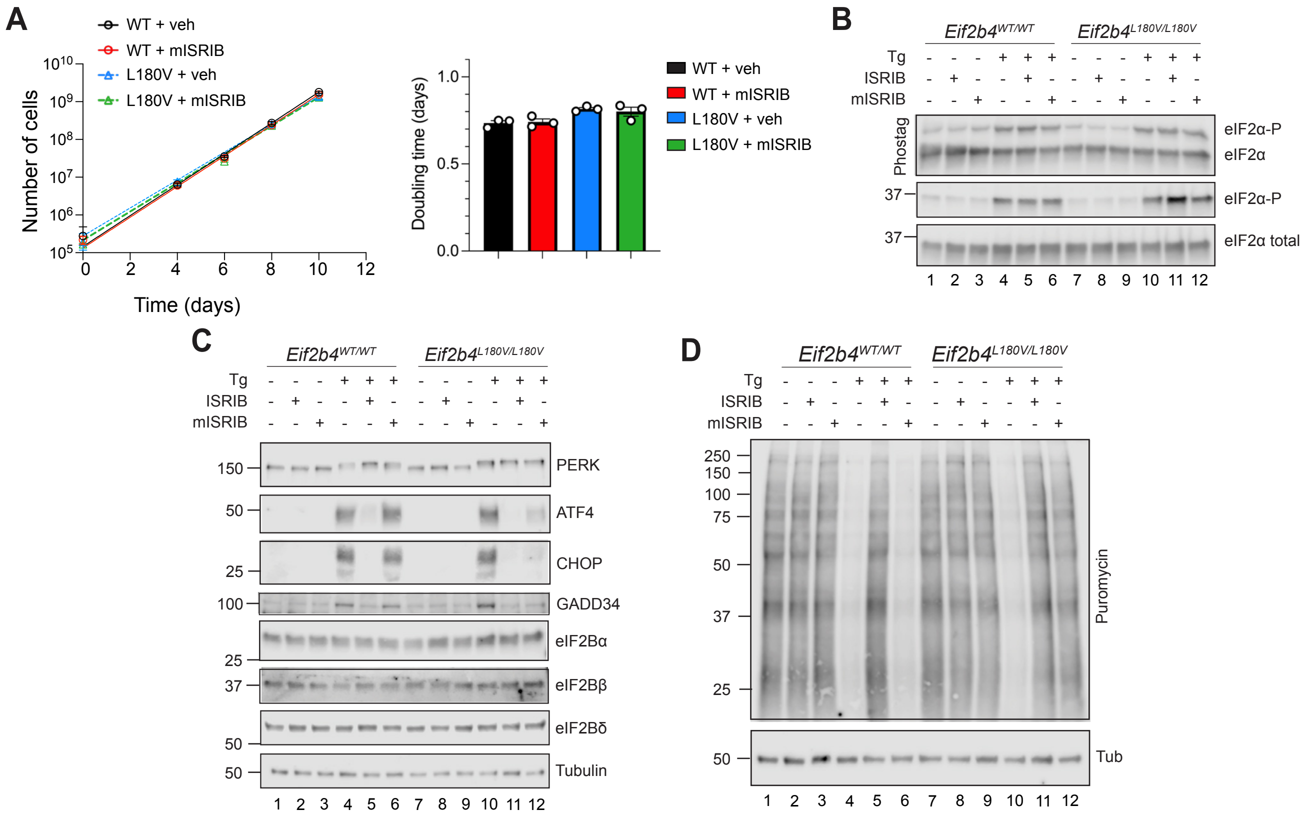
A
WT + veh
WT + mISRIB
L180V + veh
L180V + mISRIB
Number of cells
Time (days)
Doubling time (days)
B
Eif2b4WT/WT
Eif2b4L180V/L180V
Tg
ISRIB
mISRIB
Phostag
eIF2α-P
eIF2α
eIF2α total
C
PERK
ATF4
CHOP
GADD34
eIF2Bα
eIF2Bβ
eIF2Bδ
Tubulin
D
Puromycin
Tub

**Suppl. Fig. 8**

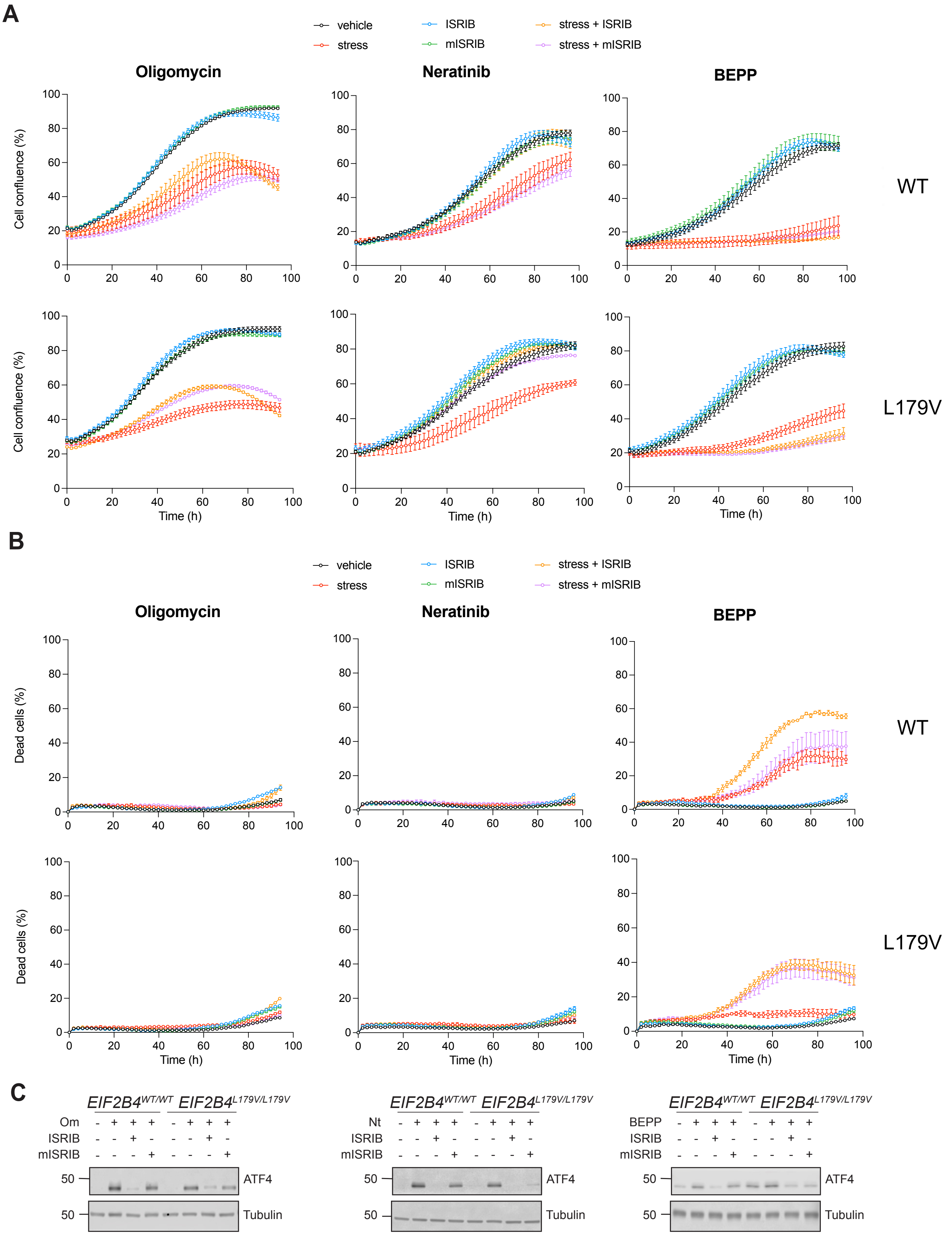
A
vehicle
stress
ISRIB
mISRIB
stress + ISRIB
stress + mISRIB
Oligomycin
Neratinib
BEPP
Cell confluence (%)
WT
L179V
Time (h)
B
Dead cells (%)
C
EIF2B4WT/WT
EIF2B4L179V/L179V
Om
Nt
BEPP
ISRIB
mISRIB
50
ATF4
Tubulin

**Suppl. Fig. 9**

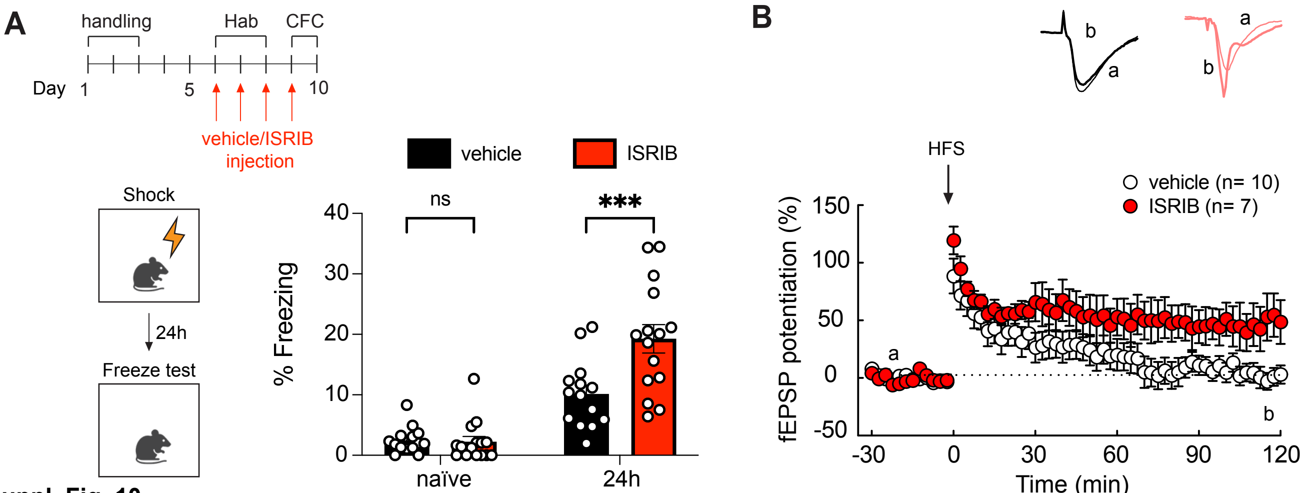
A
handling
Hab
CFC
Day 1
5
10
vehicle/ISRIB
injection
Shock
24h
Freeze test
vehicle
ISRIB
ns
***
% Freezing
40
30
20
10
0
naïve
24h
B
b
a
a
b
HFS
vehicle (n= 10)
ISRIB (n= 7)
fEPSP potentiation (%)
150
100
50
0
-50
a
b
-30
0
30
60
90
120
Time (min)

**Suppl. Fig. 10**

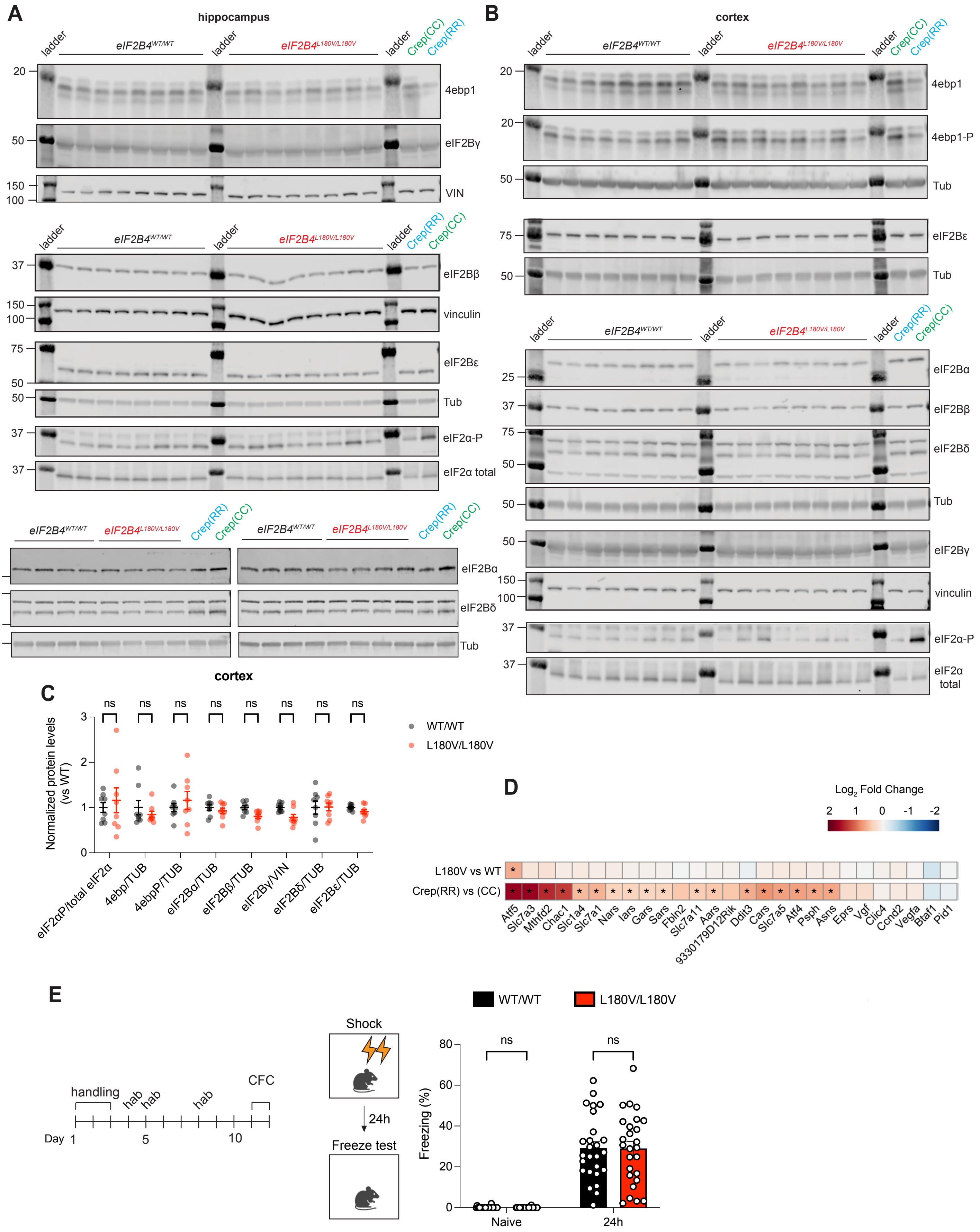

A
hippocampus
ladder
eIF2B4WT/WT
eIF2B4L180V/L180V
Crep(CC)
Crep(RR)
4ebp1
eIF2Bγ
VIN
eIF2Bβ
vinculin
eIF2Bε
Tub
eIF2α-P
eIF2α total
eIF2Bα
eIF2Bδ
B
cortex
4ebp1-P
eIF2α total
C
cortex
Normalized protein levels (vs WT)
ns
WT/WT
L180V/L180V
eIF2αP/total eIF2α
4ebp/TUB
4ebpP/TUB
eIF2Bα/TUB
eIF2Bβ/TUB
eIF2Bγ/VIN
eIF2Bδ/TUB
eIF2Bε/TUB
D
Log2 Fold Change
L180V vs WT
Crep(RR) vs (CC)
Atf5
Slc7a3
Mthfd2
Chac1
Slc1a4
Slc7a1
Nars
Iars
Gars
Sars
Fbln2
Slc7a11
Aars
9330179D12Rik
Ddit3
Cars
Slc7a5
Atf4
Psph
Asns
Eprs
Vgf
Clic4
Ccnd2
Vegfa
Btaf1
Pid1
E
handling
hab
CFC
Day
Shock
24h
Freeze test
WT/WT
L180V/L180V
Freezing (%)
Naive


Suppl. Fig. 11

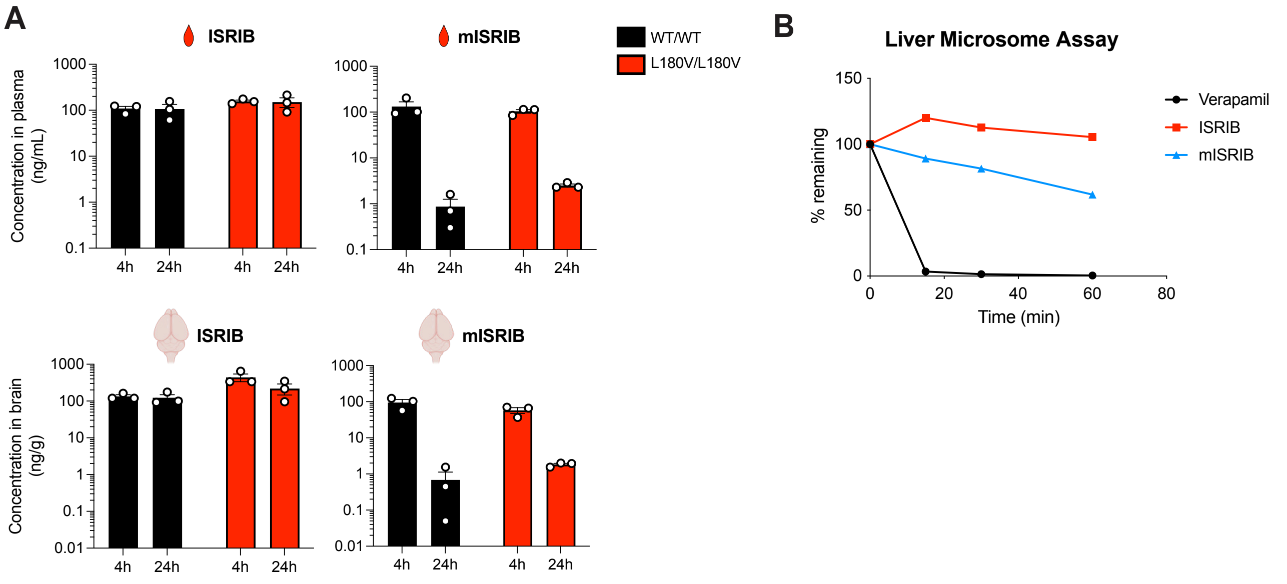
A
ISRIB
mISRIB
WT/WT
L180V/L180V
Concentration in plasma (ng/mL)
1000
100
10
1
0.1
4h
24h
4h
24h
ISRIB
mISRIB
Concentration in brain (ng/g)
0.01
B
Liver Microsome Assay
% remaining
150
100
50
0
Time (min)
0
20
40
60
80
Verapamil
ISRIB
mISRIB

**Suppl. Fig. 12**
